# Choragraph: A deep-learning approach for the analysis of spatial proteomics reveals subcellular Arabidopsis protein trafficking routes and multi-residency

**DOI:** 10.64898/2026.06.27.734956

**Authors:** Harriet T. Parsons, Tim J. Stevens

**Affiliations:** Quadram Institute; MRC Laboratory of Molecular Biology

## Abstract

Spatial subcellular proteomics provides key insights into subcellular organization but is frequently constrained by missing data values and an inability to robustly classify dual-localized proteins. To address these analytical bottlenecks, we introduce Choragraph, a deep-learning framework that uses an ensemble of deep neural networks incorporating whole-proteome cross-attention and Bayesian variational inference (BVI). Choragraph has two concurrent aims: it provides context-dependent reconstruction of missing proteomic values and predicts proteins’ subcellular compartment in a manner natively aware of multi-localisation. We applied Choragraph to a comprehensive *Arabidopsis thaliana* hyperLOPIT dataset containing 84 fractions across eight replicate LOPIT experiments. By successfully reconstructing profiles with up to 35% missing values, Choragraph incorporated over 2,000 low-abundance proteins that traditional methods would exclude. The models confidently classified 87% to 94% of singly-localised data-sufficient proteins across 14 subcellular compartments with a macro F1 score of 0.917, outperforming conventional classifiers. Crucially, Choragraph also identified over 1,000 dual-localized proteins, mapping continuous trafficking trails along the secretory pathway, highlighting functional zonation and membrane contact sites. To ensure accessibility, all data, predictions of subcellular localisation, and interactive 2D UMAP visualizations are available via an installation-free web application at choragraph.org. This framework provides a high-resolution, user-friendly resource that advances research capacity to explore subcellular location and dynamics

## Introduction

The field of spatial subcellular proteomics began 20 years ago with the development of the Localization of Organelle Proteins by Isotope Tagging (LOPIT) technique, which was first applied to *Arabidopsis thaliana* (Dunkley *et al*., 2006). The idea behind this approach is to gently distribute subcellular compartments, which are largely membrane bound, across a density gradient. This partially separates the different compartments so that they may be resolved from one another. By fractionating the gradient, and subsequently labelling and quantifying the constituent proteins via mass spectrometry, high-resolution protein abundance profiles are generated along the gradient. Because proteins residing within the same subcellular compartment exhibit similar gradient profiles (See Figure 1a for a combined overview and Supp Figure S1a for compartmental average profiles), they naturally form a cluster. By leveraging prior knowledge of a subset of well-characterized marker proteins, which have been assigned to specific compartments, these clusters can generally be assigned to distinct organelles.

**Figure 1:**
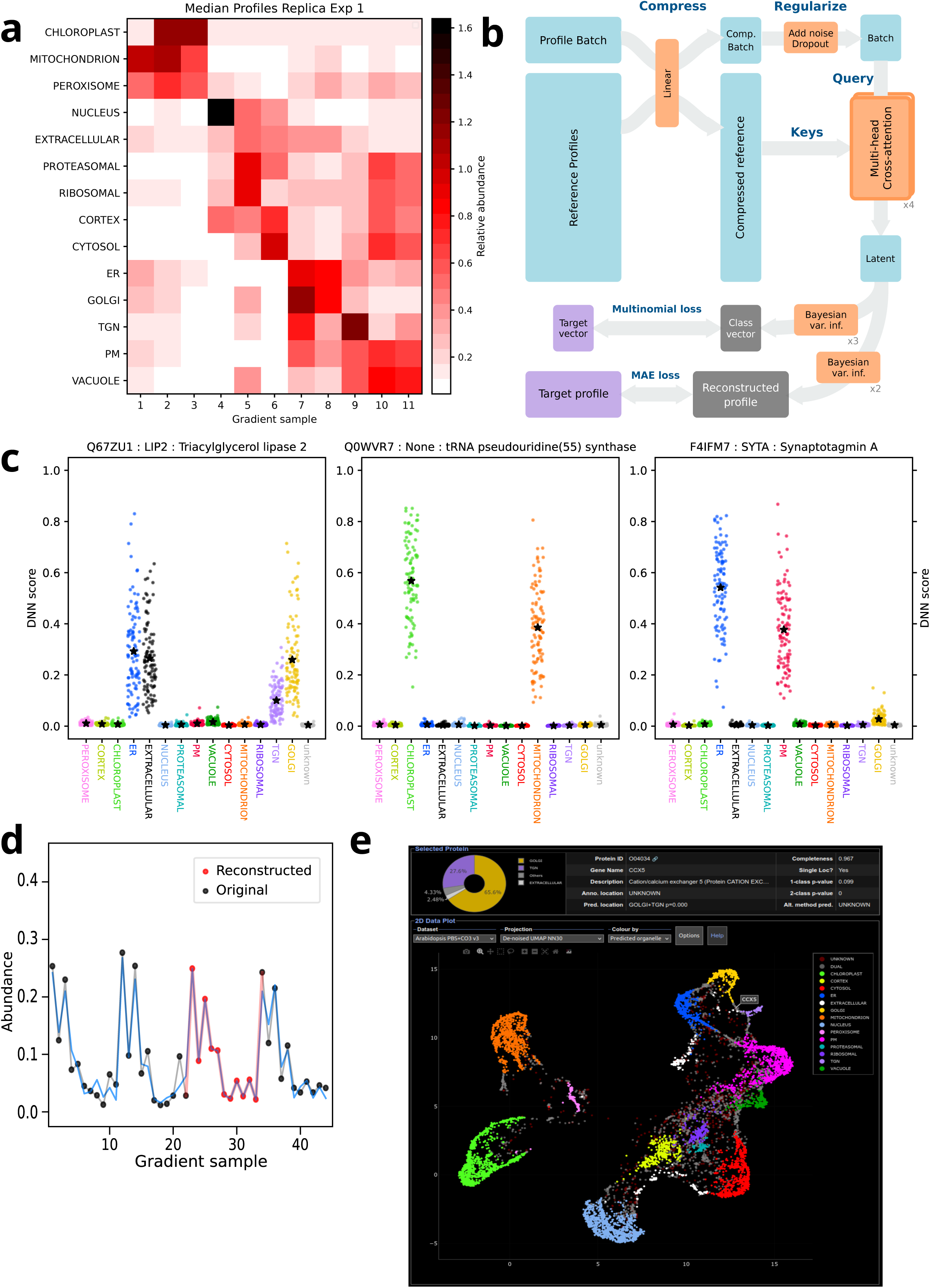
An overview of the Choragraph DNN, its inputs outputs and web application. a) A heat map of the median proteomic abundance profiles for the subcellular compartment classifications used here. The data shown relates to one replicate LOPIT experiment with 11 gradient fractions. b) A schematic of the DNN architecture used to predict (potentially mixed) subcellular location and output reconstructed abundance profiles. Input profiles are compressed to a smaller latent representation, which is then augmented by comparison to a large, proteome-wide set of latent profiles using cross attention. The final latent vector is then used for both prediction of a subcellular class vector and reconstruction of an abundance profile. c) Example DNN output scores for a selection of multi-localised proteins, illustrating the ensemble of scores that are used to estimate class proportions and corresponding classification p-values. d) An example of missing value reconstruction by the DNN. Here the red points represent where values missing in the input data have been reconstructed. e) A screen shot of the interactive Choragraph web application presenting results as an interactive 2D UMAP display.

A primary advantage of LOPIT over traditional organelle isolation methods is its capacity to distinguish resident and transient proteins within a given compartment. For example, while proteins that transiently traffic through the secretory pathway may yield a partial, minor secretory signal, the predominant mass spectrometric signal will originate from the final destination, where it is most abundant. Consequently, the protein will tend to cluster with other resident components of its destination organelle.

LOPIT and similar methodologies (Christoforou *et al*., 2016; Geladaki *et al*., 2019; Itzhak, Schessner and Borner, 2019; Schessner *et al*., 2023; McCaskie *et al*., 2025) have continuously evolved, reflecting concurrent advancements in experimental design, mass spectrometry instrumentation and chemical reagents. However, some limitations have persisted in data analysis, which have prevented the experimental datasets from being fully exploited.

One notable challenge is the need for high-accuracy imputation of missing values in protein profiles. Traditionally, missing values for protein abundance in any one of the labelled gradient fractions resulted in failed calculations for that protein and triggered its complete exclusion from downstream analysis. A nearest-neighbour based imputation method has been built into the MSnbase package at Bioconductor (Gentleman *et al*., 2004). However, the recommended limit here is 20% missing values (Hutchings *et al*., 2023), which still excludes many low-abundance proteins and discriminates against highly resolved gradient separations, where certain protein abundances can be either very high or low at either end of the gradient. Also, where a high proportion of proteins are reconstructed, with up to 20% of missing values filled, the results have not, in our experience, looked optimal.

A second, common analytical bottleneck is the inability to robustly identify proteins with gradient distributions that result from being localised to two or more different organelles. This has been, in part, addressed by the TAGM method (Crook *et al*., 2018) but stops short of directly classifying protein profiles as being best explained by specific combinations of compartments. Similarly, current frameworks lack the nuance to classify proteins that have robust, highly reproducible gradient profiles, but which do not fit either identifiable standard single- or dual-localisation patterns.

Finally, the broader utility of spatial proteomics datasets to the scientific community remains constrained by impediments to accessibility. Historically, navigating such data has required end-users to install specialized software packages, execute local scripts, and/or manually incorporate large supplementary data tables (Gatto *et al*., 2014; Breckels *et al*., 2018). To present spatial proteomics data into a genuinely accessible resource for the wider scientific community, irrespective of computational proficiency, there is a clear need for installation-free, interactive web applications. Such platforms must be easily deployable via standard web browsers through a direct hyperlink, featuring intuitive query mechanisms, dynamic data presentation, and data export functionalities, all to support typical biological workflows.

### Choragraph

Here we present Choragraph, a new deep-learning based approach to the spatial mapping and analysis of subcellular proteomics, which addresses the mentioned challenges in a single method with an accompanying web application. Choragraph uses an ensemble of deep neural network (DNN) models that employ whole-proteome based cross attention and Bayesian variational inference (BVI). This provides context-dependent reconstruction of missing proteomic values, and prediction of a protein’s subcellular localisation in a manner that is innately aware of multi-localisation.

The prediction of protein subcellular localisation by classifying proteomic profiles from LOPIT data is not an especially complex task for current machine learning approaches. The size of the data vectors (typically 2-4 replicates of 10plex or 11plex data) is tiny compared to the kinds of input used in large deep-learning models. Accordingly, with sufficiently accurate and numerous training class labels, the subcellular classification task is readily achieved with established machine learning models like SVM (Cortes and Vapnik, 1995) and EM-GMM (Dempster, Laird and Rubin, 1977).

However, here we use DNNs to move beyond a simple classification based model, to a dual purpose model that both reconstructs missing values in the data (e.g. helping to incorporate low abundance proteins) and that also classifies subcellular localisation to allow for a continuum of mixed class identities. For the latter task we especially seek dual-localised proteins with experimental profiles that can be explained as a weighted combination of two pure organelle-typical profiles. For this model we use input data (hyperLOPIT proteomic profiles) that have a singular location but train on fractional mixtures of these, to predict the original input localisation classes and their relative proportions.

Accordingly, we have created a DNN architecture, illustrated by the scheme shown in Figure 1b, to predict organellar proportions and reconstruct partially masked input profiles at the same time. Specifically, we employ a cross-attention based whole-proteome method to allow classifications and reconstructions to be based upon a context of similar proteins/profiles within the overall dataset. For the prediction of subcellular localisation, the stochastic output from the models’ BVI layers provides a mechanism to sample a distribution of possible predictions, rather than simply seeking optima. This innately leads to measures of predictive confidence, and allows the calculation of p-values to assess single- and dual-localisation hypotheses for every query protein. A typical distribution of DNN score outputs is illustrated in Figure 1c for a few example multi-localised proteins.

As shown in Figure 1d, taking a partially masked abundance profile as input our DNN generates a complete, denoised output profile that also has any experimentally missing values filled-in. Compared to typical methods for reconstructing missing values (e.g. KNN imputation) our approach allows for complex non-linear dependencies to be exploited. These may come from both intra-query replicate information and from query-dependent proteomic context, to reconstruct a significant proportion of missing values. With this, we aim to maximise the number of proteins for which we can predict subcellular localisation. Also, using the reconstruction of masked values as an objective mitigates against overtraining, as it forces the DNN model to learn the structure of the data, with an internal latent representation that maps in a continuum to the original proteomic profiles, and not to an over-optimised classification map.

Our results are available for exploration at the web app: choragraph.org, which we illustrate in Figure 1e. The Arabidopsis data used here is a combination of four replicate datasets presented in (Parsons *et al*., 2019) and four previously unpublished replicates from the same cell line. For four of the eight replicates, non-specific protein interactions have been eliminated by adding a CO₃ wash at the final centrifugation stage, after density centrifugation and gradient fractionation. Loss of functional interactions and membrane architecture was minimised by combining these with the four unwashed replicates. This gave a concatenated hyperLOPIT set of 84 fractions, which has yielded a very high-resolution subcellular spatial proteome for Arabidopsis. The capacity to usefully reconstruct gradient profiles with up to 35% missing values has added over 2,000 proteins compared to the dataset in (Parsons *et al*., 2019). With this we aim to add an informative, user-friendly resource to the research community that advances testable hypotheses relating to protein function and subcellular organisation.

## Results and Discussion

### Profile reconstruction

Table 1 shows the extent of missing values when the four unwashed replicates (unpublished) were combined with the four published CO₃-washed replicates from Parsons et al. 2019. The proportion of data loss for the two preparation conditions is around 23-24%, and this rises to 32% when all replicates are concatenated to the maximal hyperLOPIT set. As described in Methods, all data-sufficient proteins (≥ 65% complete profiles) were used in the DNN training for profile reconstruction; denoising and predicting missing values.

**Table 1:**
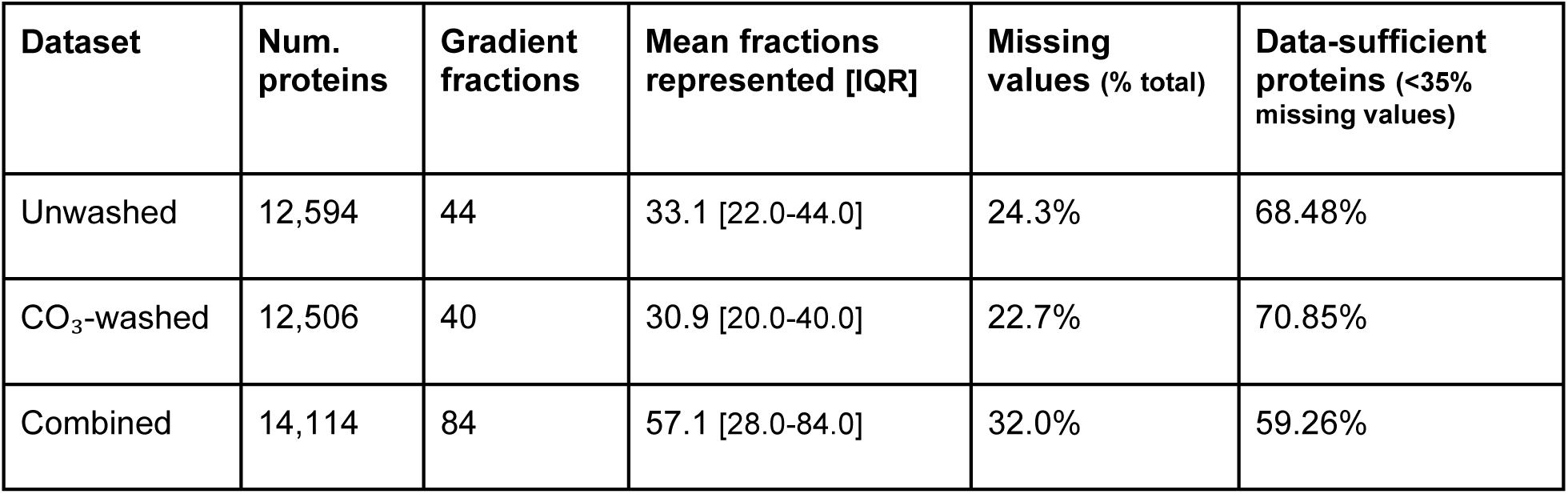
An overview of the prevalence of missing values in the LOPIT datasets studied here.

| Dataset | Num. proteins | Gradient fractions | Mean fractions represented [IQR] | Missing values (% total) | Data-sufficient proteins (<35% missing values) |
| --- | --- | --- | --- | --- | --- |
| Unwashed | 12,594 | 44 | 33.1 [22.0-44.0] | 24.3% | 68.48% |
| CO<sub>3</sub>-washed | 12,506 | 40 | 30.9 [20.0-40.0] | 22.7% | 70.85% |
| Combined | 14,114 | 84 | 57.1 [28.0-84.0] | 32.0% | 59.26% |

As shown in Figure 2a, the output of reconstructed abundance profiles (25% of which are always masked in the input during training) allows for a sensible reconstruction of missing data values. When there are a few values missing these generally correspond to low intensity values, as would be expected given stochasticity and the limits of proteomic sensitivity. When there are large numbers of points missing these often correspond to a protein being entirely missed by one replicate experiment. Nonetheless, missing values are reconstructed by the DNN in a manner that is aware of the correlations that naturally occur between replicate datasets. The overall effect of the reconstructed output is shown by the UMAP projections in Figure 2b, for proteins where up to 35% of the abundance profile may be missing. Here the reconstruction has allowed virtually all protein profiles that are initially dissimilar to the main (organelle grouped) clusters to be incorporated into the natural data structure. Visualisation of the internal, latent data representation from the DNN as a UMAP (Supp Figure S1b) shows that a large majority of the protein profiles are incorporated into the organellar clusters at an intermediate stage, before the full abundance profiles are reconstructed during the latter stages of the DNN.

**Figure 2:**
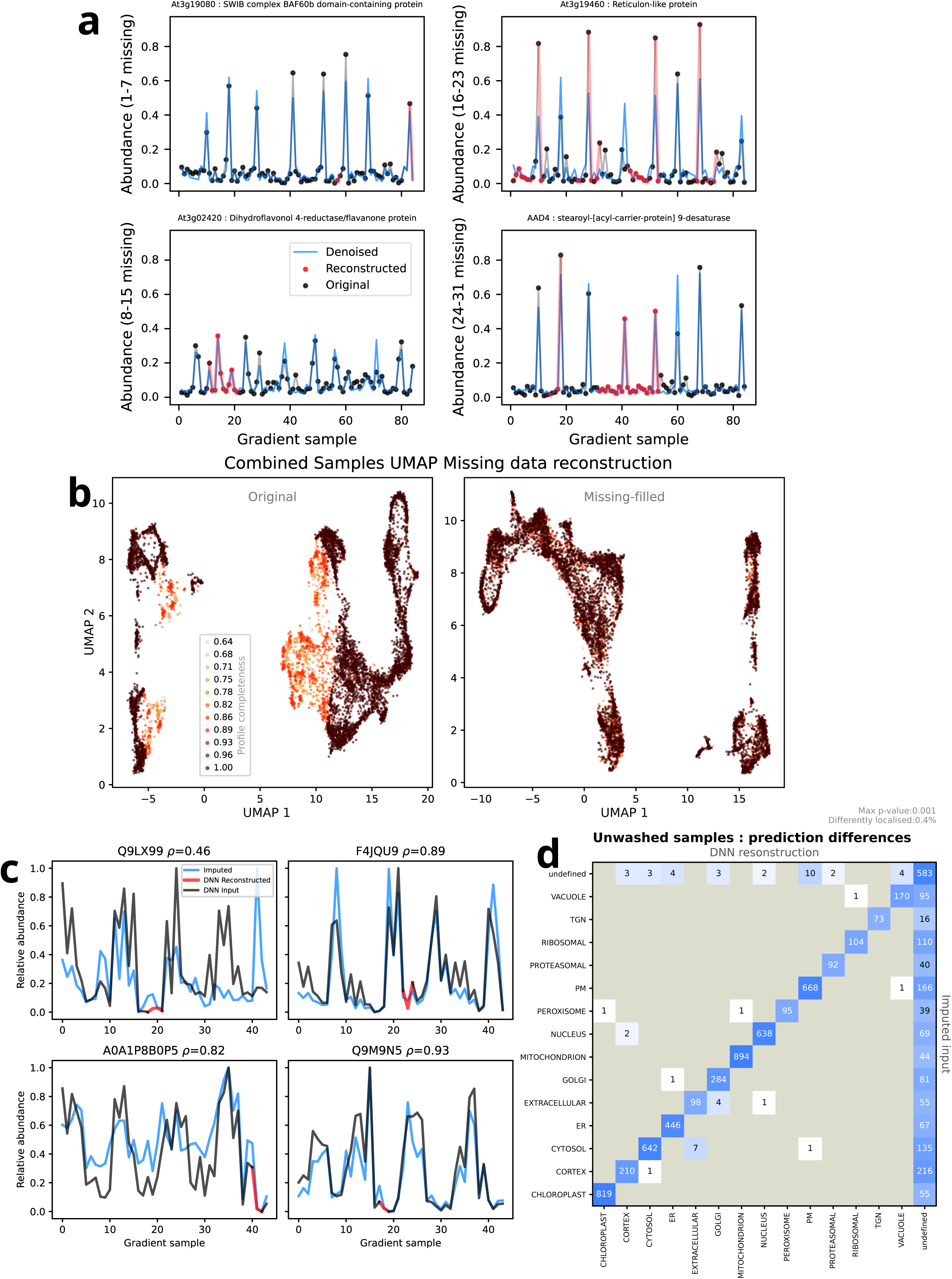
An overview of using the Choragraph DNN for reconstruction of values missing in protein abundance profiles. a) A random selection of reconstructed proteomic profiles from the Arabidopsis dataset, stratified according to the degree of missing data. The original data, with missing values, is shown in black. The full denoised profiles, which are median values from the sampled DNN model output, are shown as a blue line. The values which are reconstructed to fill-in missing values are shown in red. b) 2D UMAP projections of (left) the original, input proteomic profiles where zeros are used for missing values and (right) the reconstructed, missing-filled profiles from the DNN output. Each point represents a different protein and is coloured according to the fraction of values missing in the original profile. The data shown is for the combined dataset; eight replicate LOPIT sets. c) Comparison between KNN imputation of the data (as described at (Troyanskaya *et al*., 2001)) with a few missing values and DNN reconstruction of the same missing values. Data is shown for the first replicate of the unwashed LOPIT set. d) Comparison between single-class compartmental predictions when using the DNN to classify from the KNN imputed data and using the original data (with missing values) and reconstructing values on-the-fly. Similar to a confusion matrix, the values tally how the classifications made by one method are classified by the other method. Diagonal elements indicate identical classification and off-diagonal elements indicate differences.

We compared our DNN-based profile reconstruction method with a k-nearest neighbour (KNN) based imputation method used previously for LOPIT data (Gatto *et al*., 2014; Breckels *et al*., 2018), using recommended parameters to fill-in two or fewer missing values per replicate profile. Here we performed the KNN imputation on the unwashed dataset prior to input to the same DNN, so there were no initially missing values. Our DNN-based reconstruction and the KNN imputation both-fill in missing values in a sensible manner (see Figure 2c for a variety of profile examples, and Supp Fig S1c for DNN scoring changes) and are largely adding low-intensity values (Supp Figure S1d), though it is evident that the KNN imputation pipeline scales data columns differently. In terms of predicting single-class subcellular localisation (as described below) the pre-imputation of the data didn’t make much difference, when a classification was made (Figure 2d). However, the number of proteins for which a prediction could be made is far greater when using the DNN to reconstruct missing values. For example, with the unwashed dataset, pre-imputation led to the classification of 6,491 proteins, whereas the DNN reconstruction gave 7,860; 21% more.

### Prediction of subcellular compartment

As described in the Methods and illustrated in Supp Figure 2a, the proteins of known subcellular location, that could be used to train the DNN, were culled to remove a small fraction of profiles that were peripheral to the mainstay of their class (Wilson, 1972). This left around 2,400 proteins for each dataset, covering 14 subcellular locations which were used in the DNN’s location training, and by the other methods used for comparison.

As shown in Table 2, confident predictions were made each time for over five thousand proteins, based on the classified (marker) training sets, resulting in 87-94% of proteins with sufficient data having one or more classified subcellular locations. Notably, there are fewer borderline predictions (0.001 < p < 0.1; see Methods for p-value estimation) than unreliable ones, hinting that the single-or-dual localisation hypothesis we use is insufficient for a small fraction of proteins with more complex localisations.

**Table 2:** An overview of the number of training markers and predictions of subcellular localisation stratified by quality, for each of the datasets. Percentages for data sufficiency relate to all detected proteins, whereas the percentages for other columns relate to data-sufficient proteins

| Dataset | Proteins detected | Data deficient | Data sufficient | Classified proteins | Training proteins | Solid; $p < 0.001$ | Borderline; $p < 0.1$ | Unreliable; $p \geq 0.1$ |
| --- | --- | --- | --- | --- | --- | --- | --- | --- |
| Combined | 14,114 | 5,821 (41.2%) | 8,293 (58.8%) | 7,809 (94.2%) | 2,422 (29.2%) | 5,386 (64.9%) | 18 (0.2%) | 466 (5.6%) |
| Unwashed | 12,594 | 3,994 (31.7%) | 8,600 (68.3%) | 7,860 (91.4%) | 2,415 (28.1%) | 5,444 (63.3%) | 32 (0.4%) | 708 (8.2%) |
| CO<sub>3</sub>-washed | 12,506 | 3,641 (29.1%) | 8,865 (70.9%) | 7,767 (87.6%) | 2,410 (27.2%) | 5,357 (60.4%) | 31 (0.3%) | 1,067 (12.0%) |

The proportions of proteins with a training classification or a subcellular location prediction are also shown in Figure 3a, for the unwashed sample conditions, and in Supp Figure S2b for the other datasets. Here it is notable that the singly-localised proteins almost entirely fall within the most confident predictions, and as prediction confidence declines the proportion of proteins that cannot be assigned to either a single- or dual- localisation increases.

**Figure 3:**
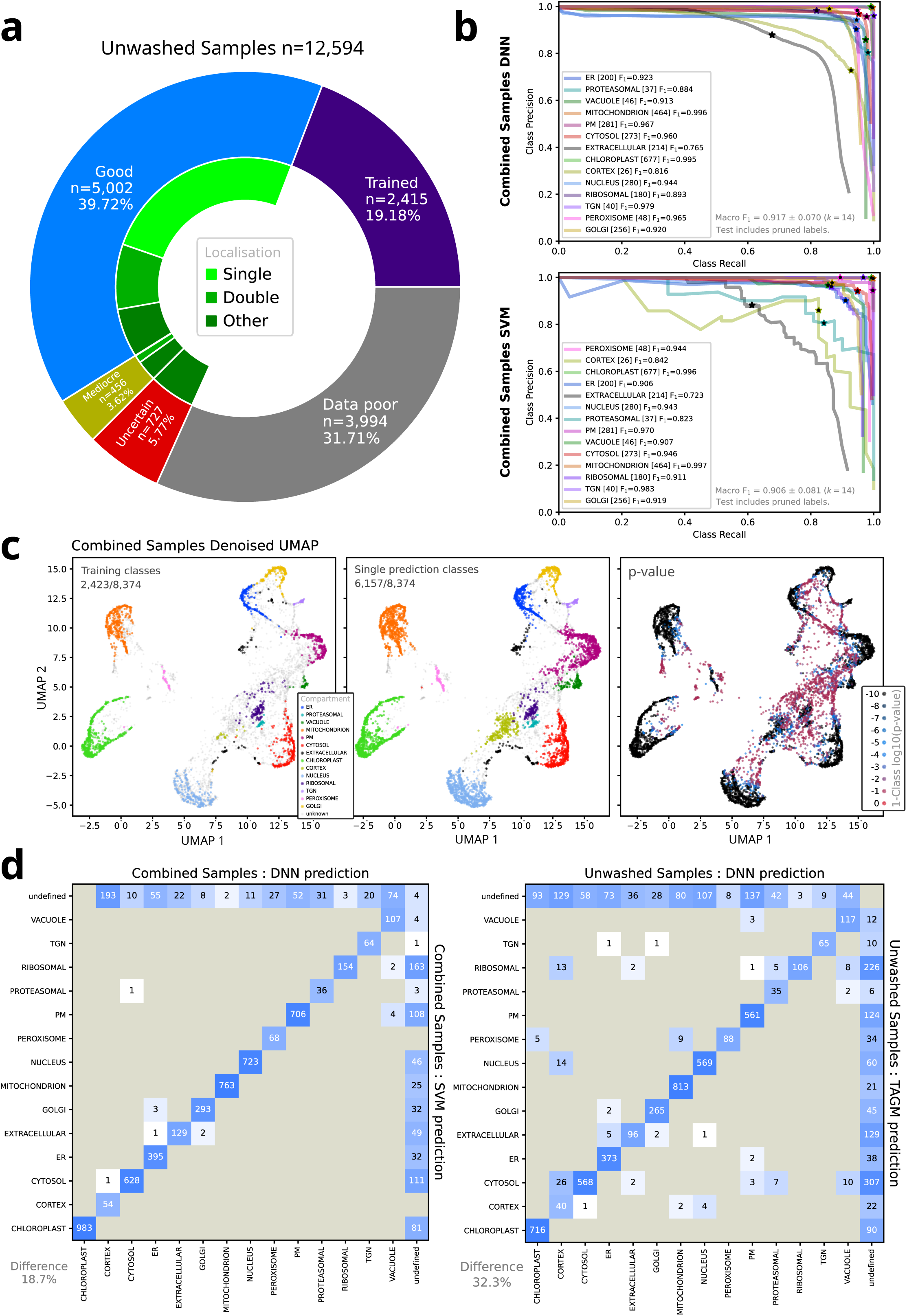
Analyses of the single-location subcellular predictions made by the Choragraph DNN. a) Overview of how protein abundance profiles were used with the DNN classification, for both the unwashed dataset. The outer ring illustrates the proportions of profiles used as training markers, excluded from classification due to missing values (>35%) or classified with various degrees of confidence, as given by p-value. The inner green ring shows the proportions of the classified profiles that were considered, single-, dual- or multi-localised. b) Precision-recall curves and maximum F1 scores for each of the singly classified subcellular location classes using DNN models (left) or conventional SVM models (right) trained on the combined dataset. Data shown are averages of ten independent test-train data splits, and the same data splits were used for both DNN and SVM. In all cases the test proteins also included markers that were pruned, and never used in training. c) 2D UMAP projections of denoised proteomic profiles coloured according to training marker classification (left), predicted single-class classification, with p1-value < 0.001 (middle) and the p1-value themselves (right). d) Comparison matrices to illustrate differences in protein subcellular classification when using the DNN compared to an SVM (left) or the DNN compared to TAGM (right) (Crook *et al*., 2018). Similar to a confusion matrix, the values tally how the classifications made by one method are classified by the other method. Diagonal elements indicate identical classification and off-diagonal elements indicate differences. Classification thresholds for the different methods were: DNN p1 < 0.001; SVM probability >= 0.8; TAGM MCMC probability >= 0.9999.

As illustrated in Table 3, the datasets have confident predictions of a single subcellular location for about 65% of the input proteins. For prediction of dual-localised proteins (see next section) the combined dataset makes more predictions and has fewer indeterminate assignments; where location is not resolved to better than three possibilities. This is likely because more replicate data increases the fidelity of discrimination, and hints that while some proteins may be genuinely highly promiscuous in subcellular location, indeterminate localisations can often be resolved with more data.

**Table 3:** The breakdown of the most confident localisation predictions (p < 1e-4) according to whether they represent single, dual or indeterminate (three or more) classes.

| Dataset | Single localised | Dual localized | Indeterminate |
| --- | --- | --- | --- |
| Combined | 3,224 (64.8%) | 1,412 (28.4%) | 341 (6.9%) |
| Unwashed | 3,209 (64.2%) | 1,025 (20.5%) | 768 (15.4%) |
| CO<sub>3</sub>-washed | 3,460 (65.5%) | 1,252 (23.7%) | 570 (10.8%) |

**Table 4:** Summary of colour schemes for data points in the 2D Plot Area of the webpage (accessible at choragraph.org).

| Option | Description |
| --- | --- |
| Predicted location | Subcellular location predictions from our DNN-based supervised machine learning. |
| Organelle Pair | A dropdown menu that presents sub-menus for selecting organelle pairs. Shows intensity gradients of ‘shared-ness’ between each selected pair from red (compartment 1) to blue (compartment 2), such that proteins with equally shared profile characteristics are yellow. |
| Annotated location | Known marker annotations used for supervised learning in this study |
| Other annotation | Sub-compartment annotation, where available (Ferro et al., 2010; Duncan et al., 2011; Hooper et al., 2017), otherwise uses supervised learning markers. |
| Alt. methods | High-confidence TAGM-MCMC predictions (Crook et al., 2018) to compare with other classifications (predictions only derived from the unwashed sample data). |
| 1-class p-value | Heat gradient showing the single-class hypothesis $p_1$-value: how confident we are of each protein belonging to a single location, from most confident (blue) to least (red). |
| 2-class p-value | Heat gradient showing the dual-class hypothesis $p_2$-value: how confident we are of each protein belonging to either a single or dual location, from most confident (blue) to least (red). |
| Dualness | Heat gradient showing the degree to which the abundance profile appears to derive from two compartments rather than one. Specifically, the values are the gain in the fraction of DNN scores $> 0.8$ when combining primary and secondary classes. |
| Completeness | Heat gradient showing how complete a protein’s original data profile is, in terms of missing values, from least complete (red) to most complete (blue). |

Training of the DNN resulted in models that made very few errors when making class predictions (0.7%; see Supp Figure S2c) for the test profiles, which were excluded from each model’s training. This is not surprising, given that peripheral class markers were pruned prior to training and most class boundaries were clear under UMAP projection (see below). Of the peripheral markers that were disregarded prior to training around 29% have contrasting re-assignments upon DNN classification. These most notably include Golgi proteins being reclassified as ER or extracellular; unsurprising given secretory trafficking and cargo, and extracellular proteins having a variety of reassignments; resulting from diverse profiles as cellular cargo and relocation to the nucleus (e.g. under cellular stress)

A precision-recall analysis (Figure 3b, Supp Figure S2d), which included the peripheral markers disregarded from DNN training, shows that the DNN is an effective classifier (macro F1 = 0.917 ± 0.070), though the models are not solely designed for this purpose. Also, the DNN performs slightly better than a conventional SVM based classifier (macro F1 = 0.906 ± 0.081, Figure 3b, Supp Figure S3a). The least predictable classes are extracellular and cortex compartments; understandable given that extracellular proteins are distributed across multiple smaller profile clusters and that cortex profiles mix with ribosomal ones (see Figure 3c). Though the DNN and SVM classifications are highly similar for many proteins, the different approaches are classifying different subsets of profiles (Figure 3d). As is clear in the UMAP projections (Supp Figure S3b), the DNN and SVM models have somewhat different decision boundaries for classification.

UMAP 2D projections of the denoised profiles (Figure 3c) and latent vectors (Supp Figure S3c) show that the confident predictions of single subcellular location naturally follow where the marker labels were present in the training class, and tend to cover the extent of the profile clusters that are distinctly separate. It is also notable that singular classification has not generally spilled into unassigned regions of the UMAP and that there are clear trails of (projected) profiles that span between organellar clusters, these have high p1-values (single-class) hinting that they may be better explained as dual localised.

Comparison of DNN single-class predictions to TAGM MCMC method high-confidence predictions trained on identical data (Figure 3d), shows that the TAGM predictions are broadly similar, but there are notable further predictions for the cytosol, ribosomal and nucleus classes. Inspection of the classified UMAP projection (Supp Figure S3b) shows that these compartments, together with the ER, have somewhat more expansive assignments than with the SVM or DNN models.

### Multi-class prediction

As illustrated in Figure 4a, and as we may expect, the profiles that we identify as dual localised tend to occur between the clusters of profiles that have clear single-class identities, where the training markers are found. Analysis shows that the prediction of dual-localisation does not correlate with the degree of missing or filled-in values (Supp Figure S3d). The dual-class profiles tend to be present within the UMAPs as elongated, peripheral trails leading from, and between, single-class groups. These trails appear to derive from different proteins having a continuum of different relative compartmental proportions, at least as far as the abundance profile is concerned. Using the endoplasmic reticulum (ER) as an example (Figure 4b), we can see different trails of profiles lead separately to classifications involving the Golgi, plasma membrane, and trans-Golgi network. Similar transitional features are also visible in the maps for many of the compartments studied, as shown in Supp Figure S4.

**Figure 4:**
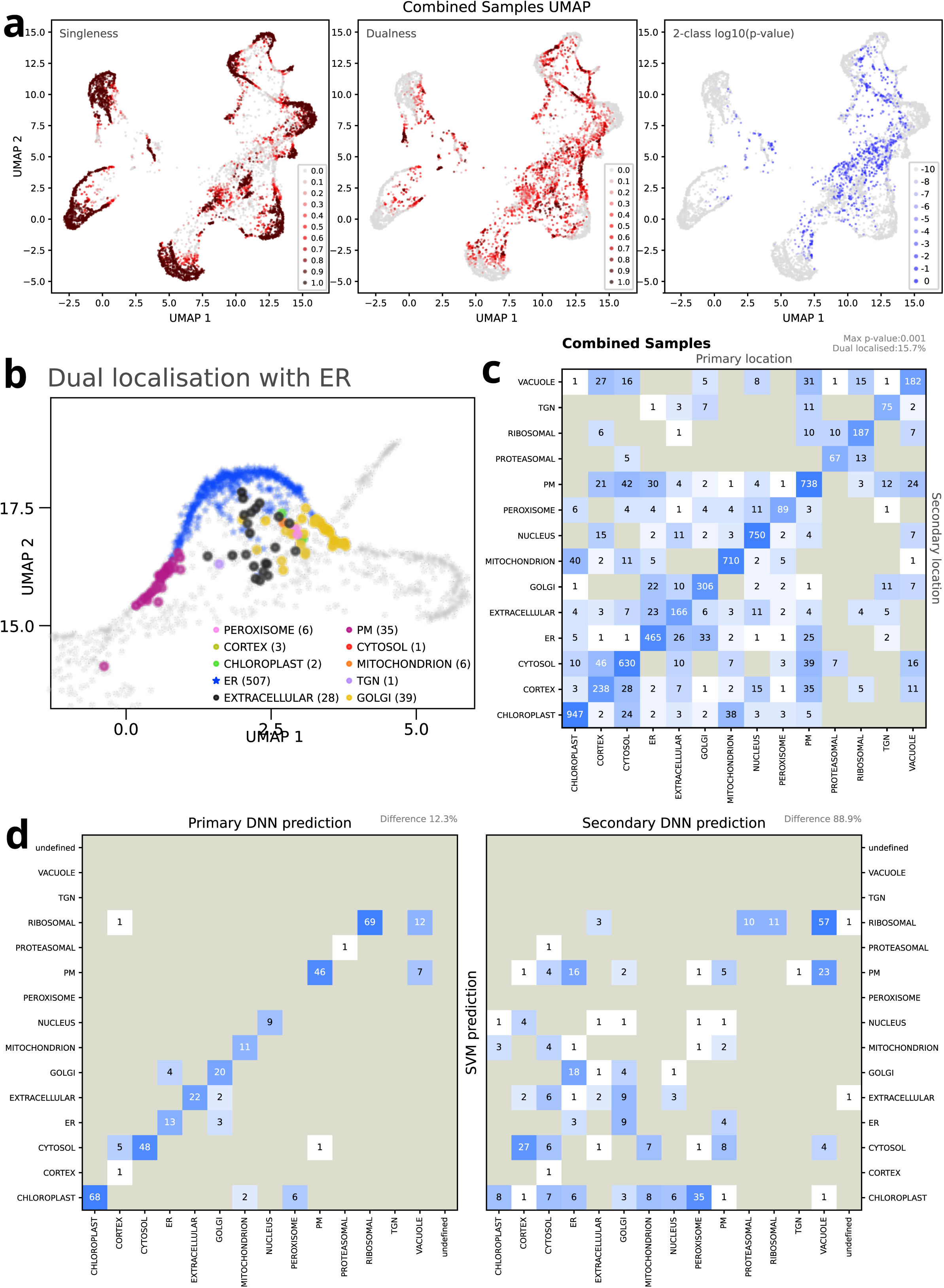
Analyses of the dual-location subcellular predictions made by the Choragraph DNN. a) 2D UMAP projections of denoised proteomic profiles coloured according to singleness; the fraction of sampled primary class scores > 0.8 (left), dualness; the gain in the fraction of scores > 0.8 when combining primary and secondary classes (middle) and dual-class p2-value (right). b) 2D UMAP projections of denoised proteomic profiles, from the unwashed dataset, with predicted single- and dual-localised endoplasmic reticulum (ER) proteins highlighted. ER-only proteins are indicated with a blue star, dual-localised proteins are indicated with circles and coloured according to the non-ER component of the classification. c) Comparison matrix to illustrate the DNN assignment of proteins to one or two subcellular classes using the combined data. Diagonal elements indicate assignment to a single-class (p1 < 0.001). Off-diagonal elements indicate assignment to dual classes (p1 > 0.001, p2 < 0.001), as specified by column (primary class) and row (secondary class). d) Comparison matrices to tally how dual-localised proteins identified by the DNN are classified by the SVM models. Diagonal elements indicate identical classification. The matrices on the left refer to the primary DNN classification (with the largest proportion) and the matrices on the right refer to the secondary DNN classification. Classification thresholds for the different methods were: DNN p1 > 0.001, p2 < 0.0001; SVM probability >= 0.8.

The overall extent of the dual classifications is shown in Figure 4c, for the combined datasets and in Supp Figure S5a for the separate wash conditions. It is clear that some compartmental pairs do not have any predicted dual localised proteins, while for others it seems common. Many proteins are dual localised along the secretory pathway (ER, Golgi, TGN, PM, Extracellular), as we may anticipate, and there is clear support for ER-PM contact sites (55 proteins). Dual mitochondrion-chloroplast proteins are especially common (78 predicted). It is notable that we do not predict any clear mixing between the nucleus and the (free) cytosol class. This occurs because our cortex compartment takes on this role; the cortex was instigated to represent the microtubule associated component of the cell interior.

When we compare the DNN dual-class predictions with the SVM (Figure 4d) and TAGM MCMC single-class predictions (Supp Figure S5b), many of the SVM or TAGM predictions do match one of the DNN dual-location options, and usually this is the primary one. For the SVM models the match to the primary DNN class is particularly good, at 88%. It is notable that TAGM predictions spread more into the DNN dual-class predictions than SVM models, especially for the compartments where the TAGM predictions are particularly extensive.

### Biological Insights

Subcellular spatial proteomics has been very successful in assigning proteins at the core of clusters to single locations but the lack of specific locational information on dual- or multi-localised proteins has limited the functional insights that can be gleaned from it. By giving locational context to every protein, even when the context is, specifically, ‘Unknown’, we gain another level of biological insight from this technique. Many, but not all, of the proteins mentioned by name in the text below are included in the accompanying figures. Readers may actively follow the next sections with the web application, changing colour schemes and filtering on protein names/properties as they go.

### Compartments and trafficking routes

One of the greatest benefits of Chorograph is its capacity to visualise gradients of changing affiliation from one location to another. Consequently, Choreograph provides insight to the routes of trafficking proteins and their protein cargo around the cell. Here, rather than describe cluster contents, we focus on the links between clusters.

#### Ribosomes and the cytoskeleton

Ribosomes form a tight cluster interspersed with various ribosome- and RNA-binding proteins. RACK proteins that attach ribosomes to the cytoskeleton (Ceci *et al*., 2012), and the motor protein myosin, are found between this and the cortex cluster. The cortex cluster itself contains proteins complexes and vesicular trafficking machinery that attaches to the cytoskeleton, such as the anaphase-promoting complex, COP9 signalosome, translation initiation complex subunits, coatomer proteins, SEC23 and SEC24, COG proteins and certain Golgins, amongst others.

#### The Endoplasmic Reticulum

Protein synthesis at the ribosome is followed by translocation into the Endoplasmic Reticulum. In the combined dataset there is evidence that membrane structures have been preserved as functional zonation can be seen within clusters. The Endoplasmic Reticulum Membrane Protein (ERMP) import complex and Sec61 subunits cluster at the far right of the ER cluster, with Protein Disulphide Isomerase-like proteins (PDILs) at the ER-curve apex (Figure 5a). Emp24-domain proteins, involved in ER-Golgi vesicular trafficking (Chen, Qi and Zheng, 2012), are in the centre. CYP71 and CYP72 proteins are found on the PM-proximal bottom of the ER curve, with Reticulons on the string of data points leading to the PM (Figure 5A). Subunits of the Oligosaccharyltransferase (OST) complex cluster on the upper curve leading to the cis-Golgi, where proteins modified by this complex will get modified yet further. The ER is a reticulated compartment of sheets and tubules, whose structure is sensitive to CO3 washing. This zonation is less distinct in the CO3-washed dataset, but similarly distinct in the unwashed dataset, suggesting a good balance between specificity and structural integrity has been achieved in the combined dataset.

**Figure 5:**
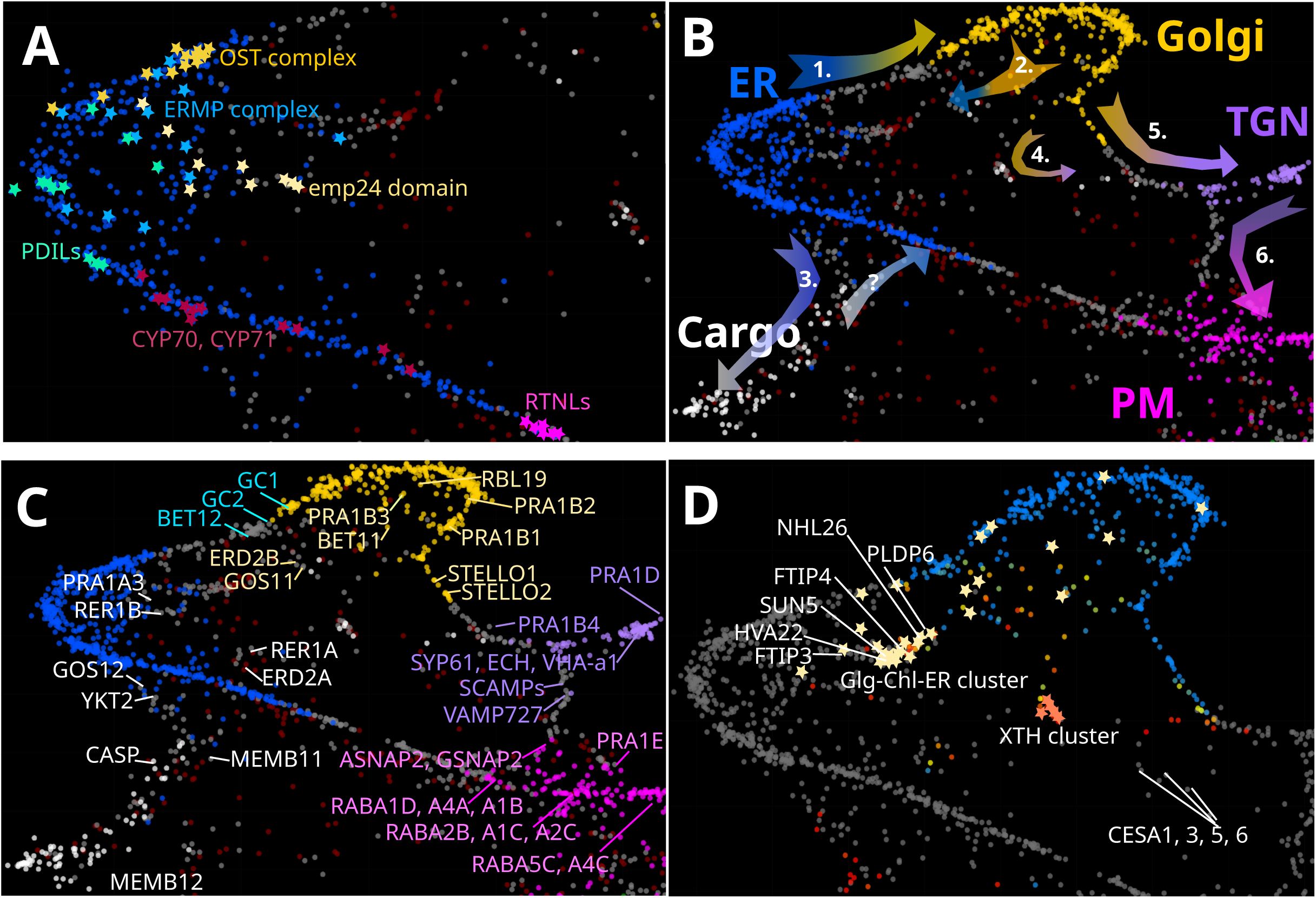
Distribution of select proteins in the early secretory system (ER, Golgi and TGN) in the Combined dataset. a) Resident endoplasmic reticulum families are highlighted with different coloured stars, showing how protein families can be resolved within the ER cluster (blue data points), and how function affects position within the cluster. Pale yellow stars denote emp24-domain, or p24, proteins. Yellow stars denote the Oligosaccharyl Transferase (OST) complex proteins, pale blue stars the endoplasmic reticulum membrane protein (ERPM) complex and turquoise stars denote Protein Disulphide Isomerases (PDILs). Plum stars show protein from Cytochrome P450 families 70 and 71. Magenta stars show reticulon-like proteins (RTNLs). b) Known and inferred directions of protein movement around this part of the secretary system. Arrow numbers are referred to in the main text. Data points are coloured using the ‘Predicted Organelle’ option in the Colour Scheme dropdown menu of the Choragraph web application. Arrows are coloured with a gradient from departure to destination compartment. c) Trafficking proteins that are considered likely candidates for managing cargo between the compartment clusters in this figure. Not all proteins mentioned in the text are labelled in this figure. Additional proteins can be found using the text and filter functions in the Protein Table area of the webpage. Gene names are coloured according to the compartment in which proteins likely function. Cyan labels denote the cis-Golgi and white labels the ER-extracellular compartment defined by CASP and MEMB11. d) Proteins involved in two putative trafficking routes within the early secretory system. Pale yellow stars denote proteins within the early secretory system with >2% chloroplast and >5% Golgi components in their density gradient profiles. Select proteins involved in plasmodesmata formation, trafficking and/or membrane curvature have been labelled. Orange stars denote Xyloglucan endotransglucosylase/hydrolase (XTH) enzymes. For context to the relative position of this XTN cluster, CESA subunits have also been labelled. The colour scheme has been set to highlight the TGN-Extracellular Cargo pair.

#### From the ER into the Golgi and back again

On the Choragraph web app, use of the colour scheme to ‘Other annotation’ in any of the datasets reveals the entrance to the Golgi by way of the cis-Golgi. Alternatively, colours can be set to highlight the ER-Golgi organelle pair, or the table can be filtered for proteins with >15% ER and >15% Golgi. The direction of protein flow, as evidenced by these three visual formats, is summarised by arrow 1 in Figure 5b. Trafficking proteins operating at the cis-Golgi include Golgin family proteins GC1, GC2, the SNARE BET12 and Golgin family proteins AT5G15880 and AT4G30090 (Figure 5c). Much of the COPI and COPII recycling machinery (Aniento *et al*., 2021) is located in the cortical cluster, as the majority of these protein populations are attached to the cytoskeleton. However, Golgi-resident trafficking machinery is still identifiable. Inside the cup-shaped Golgi cluster we see BET11, GOS11, an inactive rhomboid-like protein RBL19 (Lemberg and Adrain, 2016), SNARE-associated family protein AT1G71940, retrograde transport protein AT1G50120, PRA1B1 -B3, PRA1A3, lumen receptors ERD2B and RER1C (Figure 5c). Arrow 2 in Figure 5b captures the possible direction of retrograde transport.

CASP and MEMB11 are proteins operating at the ER/cis-Golgi interface (Chatre *et al*., 2005; Fougère *et al*., 2023). We see both of these proteins in a diffuse ER extension containing extracellular, dual-localised and unknown proteins that extends out from the ER, outside of the ER-Golgi-TGN-PM connection (arrow 3, Figure 5b). At the ER ‘start’ of this extension we see the Golgi SNAP receptor GOS12 and R-SNARE YKT61. At a possible re-entry/recycling point is a cluster of putative ER lumen receptors AT4G38791, AT1G19970, AT2G21190, RER1A and ERD2A (Figure 5c). This ER extension seems connected to later Golgi trafficking, as most proteins have 2 -12 % TGN profile composition. It is, approximately, a mirror image of the TGN (Figure 5b); in a multi-dimensional data space, they might be more similar than is apparent in the 2D UMAP projections.

#### Golgi to chloroplast trafficking

The existence of a Golgi to Chloroplast trafficking route is known, as certain chloroplastic proteins are modified in the Golgi, but few proteins have been assigned to it (Baslam *et al*., 2016). After filtering the combined dataset for proteins with >5% Golgi and >2% Chloroplast, the early half of the Golgi and the interior of the Golgi curve are labelled (Figure 5d). Modified chloroplast proteins seem therefore to be leaving before the later Golgi cisternae. Several dual-localised proteins within the chloroplast cluster have Golgi profile components, suggesting trafficking by vesicular transport instead of, or as well as, ER-Chloroplast MCSs. NHL proteins have been implicated in trafficking (Xu *et al*., 2025) and many are present in this zone, including NHL26. NHL26 has substantial chloroplast, golgi, ER and cargo profile components. It operates at plasmodesmata and its overexpression has recently been shown to alter transcripts for a chloroplastic glucose transporter (Vilaine *et al*., 2013). We do not see the same transporter in our data but see two annotated plastidic glucose transporters (AT5G16150, AT1G79820) that have strong Golgi and Golgi/Chloroplast profiles. SUN5 sits in the centre of a dense ER subcluster containing a high proportion of proteins with an ER-chloroplastic-Golgi profile composition (Figure 5d). SUN5 is, in part, responsible for the organisation of ER-derived organelles, ER bodies (Ikeda *et al*., 2026). The ER-membrane curvature protein HVA22 is also found in this structure, raising the possibility that ER bodies are trafficked to the chloroplast. Interestingly, many plasmodesmal proteins, such as FTIP3 and 4, PDLP6, MCTP and HIPP family proteins are also found in this cluster. Their profiles all contain some degree of chloroplastic contribution, even though plasmodesmata are not physically attached to chloroplasts.

#### From the Golgi to the TGN and back again

STELLO1 and 2, responsible for packaging CESA subunits into post-Golgi microtubules-associated cellulose synthase compartments (MASCs) (Zhang *et al*., 2016), seem to mark the transition from Golgi to TGN (arrow 5 Figure 5b, Figure 5c). Trafficking proteins on this route include the GTPase GB1 (AT5G52210), PRA1B4 and vesicle transport protein MDA7.6. Consistent with MASCs operating independently from the TGN (De Caroli *et al*., 2020), CESA subunits 1, 3, 5 and 6 are located close to, but not in the TGN (Figure 5d). Within the dense, TGN-resident cluster we see known TGN residents VHA-a1, SYP61, 42 and 43, ECH, VTI12, VSR1, 3, 4 and 7, PRA1D, YIP proteins and many documented (Qi *et al*., 2024) PM cargo proteins (Figure 5c). Using a colour scheme to highlight Extracellular-TGN proteins shows recycling pathways operating in the interior of the ER-Golgi-TGN-PM ring. At the centre of this is a small cluster containing most of the xyloglucan endotransglucosylase/hydrolase (XTH) enzymes found in this study, amongst other proteins, and clearly undergoing exchange with the TGN (Figure 5d, arrow 4 Figure 5b).

#### From the TGN to the PM

The exit point from the TGN can likewise be viewed by using a colour pairing for TGN and PM. Here we see familiar PM-TGN recycling cargo such as PIN1, COBL7, Callose Synthases 9, 10 and 12 and Korrigan. In the late-TGN region are located SCAMP1 – 4, RABDs, VAMP727 and TET8, amongst other trafficking proteins (Figure 5c). As this route feeds into the PM cluster, we encounter VPS20.1, VPS20.2, GSNAP2 and EXO70G1, with VAMP721, ASNAP2, SEC15A and many RABAs nearby (arrow 6 Figure 5b, Figure 5c).

#### Trafficking routes to the Vacuole

Arcing round the PM cluster towards the Vacuole, RABAs give way to RABEs and RABGs. The RABG subfamily operates on the TGN-PM-MVB-Vacole pathway. In the combined dataset, setting the paired organelle colours to TGN-Vacuole highlights this route, going across the bottom of the PM cluster. Leaving the TGN, after the TGN-vacuolar sorting SNARE VAMP727 (Ebine *et al*., 2008), is a string of turquoise-yellow points marking the TGN -PM/Vacuole junction. Along this trail we find known trafficking agents in the TGN-MVB pathway such as PRA1E, VPS55, CHMP1A. We also find vacuolar vesicle trafficking protein GRV2 and exocyst subunits SEC5A, SEC5B, SEC8 and RABGs (Figure 6a). A thick stream of VHA subunits, which are trafficked along this pathway, leads to the apex of the vacuolar cluster (red stars, Figure 6a). Here the v-SNARE VTI11, SYP22 and VPS39 are located, for the docking of MVB vesicles to the tonoplast membrane. VPS9A,11,18 and 33 seem positioned to act at the cytosolic entry point to the vacuole (Figure 6b). The cytosolic-vacuole junction also marks the far end of the PM cluster, around which we see VPS36, a ring of ISTL and TOL proteins (cyan stars, red stars Figure 6b), indicating the entry point from endocytosis into the MVB. An ER autophagy pathway may also be entering the vacuole, as a line of ATG8 proteins (Aniento *et al*., 2021) extends from the PM-edge of the ER cluster to go towards VAMP724 at the vacuole periphery (green stars Figure 6b). A thin string of orange -turquoise proteins highlighted when the colour scheme is set to Vacuole-Peroxisome end at Myosin XIG at the vacuolar periphery, indicating yet one more possible route into the vacuolar cluster (not shown in Figure 6b).

**Figure 6:**
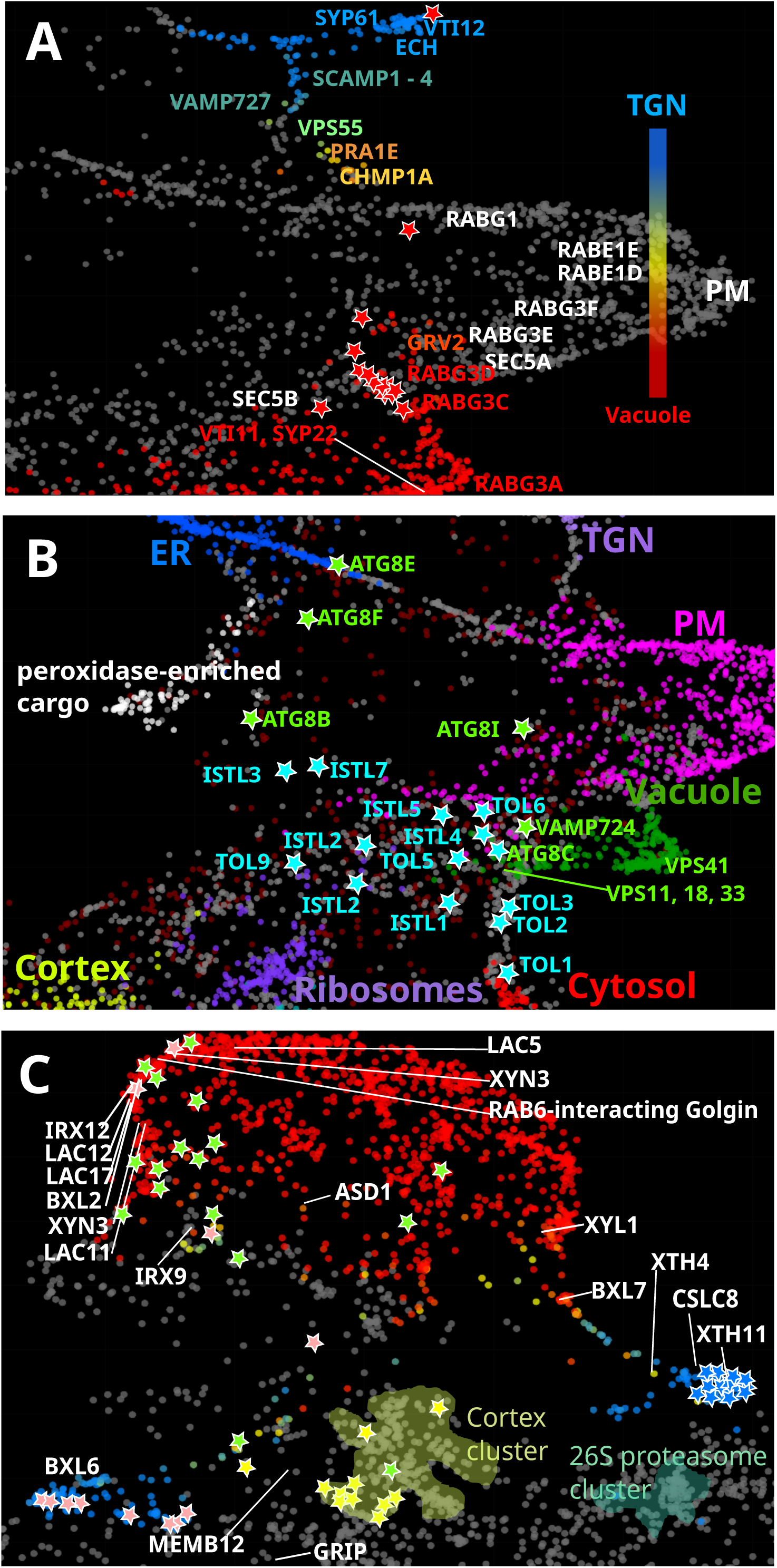
Distribution of select proteins in the late secretory system (TGN, PM, Vacuole and extracellular cargo compartments) in the combined and unwashed datasets. a) In the combined dataset the colour scheme has been set to highlight proteins localising between the TGN and Vacuole, effectively colouring the TGN-MVB-Vacuole trafficking route. Select proteins of interest have been labelled, but not all proteins named in the relevant paragraph of the results. Label colouring reflects the proportional mix of TGN and vacuolar characteristics, as shown in the colour bar. Proteins labelled in white were not highlighted by this pairing, but still contain relevant profile traits. Red stars denote subunits of the Vacuolar H⁺-ATPase. b) Proteins involved in endosomal sorting pathways to the multi-vesicular body (MVB) and degradative trafficking pathway to the vacuole, with labels coloured by protein pathway. Green stars and labels refer to the autophagy pathway. Cyan stars and labels to the endocytosis and endosomal sorting pathways. Data is from the combined dataset and data points are coloured using the ‘predicted organelle’ option. c) Part of the unwashed dataset, with the colour scheme set to highlight Nuclear-Extracellular proteins. Peroxidases are denoted by pink stars; the peroxidase-enriched vesicle referred to in the text is at the bottom left. Expansins and RALFs are marked by blue stars; the expansin/RALF vesicle is at the bottom right. Proteins likely to be involved in trafficking the vesicle are marked: yellow stars represent COG and Golgin proteins, and GRIP and MEMB12 are labelled. For context, the ‘cortex’ and 26S proteasome cluster positions are shown. Motor proteins are marked as green stars. Proteins involved in xylan biosynthesis are labelled, along with a Golgin of particular relevance.

#### Extracellular cargo

The extracellular space is not represented in our datasets as experiments were conducted on cell suspension cultures and the cell wall was digested away prior to cell rupture. What extracellular proteins have been detected are, therefore, present as cargo. Including these as a marker set gives us a guide to the presence of vesicles and certain trafficking pathways. It must be remembered, however, that these proteins will only guide us to vesicles and routes for extracellular cargo. Other types of cargo remain largely invisible.

Amongst the extracellular proteins, three functionally related clusters stand out: the XTH secretory cluster mentioned earlier, a cluster of expansins, alkalisation factors and cell-wall loosening enzymes between the cytosol and the nucleus, and a region enriched in peroxidases at the tip of the MEMB11-ER/cis-Golgi extension (Figures 5c, 6b).

Seen most clearly In the unwashed dataset, the peroxidase- and expansin-enriched clusters (pink stars and blue stars in Figure 6c, respectively) are proximal to golgins, GRIP, the COG complex and other trafficking machinery. This includes Trans-Golgi network-localized SYP41-interacting protein 1 (TNO1), confirming interaction with the TGN (Roy and Bassham, 2017), despite separation in the 2D projection (Figure 6c). Here, MEMB12 has repositioned away from the ER, as it was in Figure 5c, staying with the peroxidase-enriched cluster.

VSR6 helps traffic proteins in the late secretory pathway (Zouhar, Muñoz and Rojo, 2010). We see VSR6 at the nuclear periphery, along with a C1 domain protein that is also likely involved in protein trafficking. IRX9 appears at the Nucleus periphery in the buffer-washed dataset (Figure 6c), joined by IRX7, 10 and 15L and CESA4 in the combined dataset. Notably, even in the unwashed dataset, IRX7, 10 and 15L exhibit a decidedly peripheral Golgi localisation. It seems highly curious at first that cell wall proteins, particularly xylan biosynthesis enzymes, should localise here. However, around the proteins highlighted in the Extracellular -Nuclear pairing are members of the nuclear envelope LINC complex and many kinesin proteins (Green stars, Figure 6c). Again, this is most clearly seen in the buffer-washed dataset, suggesting that vesicles are preferentially interacting with microtubules attached to the LINC complex (Zhong *et al*., 2024; Cai *et al*., 2025). This suggests that the Golgi and post-Golgi trafficking of IRX7, 9, 10 and 15L is subtly different from that of IRX9H, 10L and 14, which are undoubtedly Golgi-localised in both datasets. This is, in part, consistent with previous observations (Brown *et al*., 2011). In both datasets, IRX14H localises to the MEMB11-ER-cis-Golgi cargo recycling loop identified above.

IRX12 and other laccases involved in lignin polymerisation, as well as the xylosidase BXL2, all locate deep within the nuclear cluster. This distribution also suggests involvement of microtubule attachments, likely by a vesicle as a RAB6-interacting Golgin is nearby (Figure 6c). Possibly the entirety of these protein populations are interacting with this Golgin, so there is no remaining population to equilibrate them away from the nucleus

#### PM to the Nucleus

Whilst Golgi and extracellular proteins at the nucleus are most likely a consequence of positioning on the cytoskeleton during cell rupture, a trafficking route between the PM and Nucleus is detectable. Using colour pairing for PM-Nucleus and filtering for proteins with >10% Nucleus and >10% PM profile composition effectively highlights this route. Along we find heat and cold shock proteins, consistent with abiotic stress signals being transmitted to the nucleus this way. Tubulin, Kinesins, chaperone proteins, including the nuclear-cytoplasmic WPP1, and subunits of the Chaperonin T-complex and ESCRT subunits VPS2.1, 2.3 and 36 are present. Vesicle unloading evidently occurs at the nuclear end of this route as we see RANGAP1 and 2, which have distinctly nucleus-PM-cortex profiles. Several nuclear pore proteins are present, as well as KAKU4, an inner nuclear envelope protein involved in transmitting stress signals to epigenetic modifying proteins.

#### Membrane contact sites and dual targeted proteins

The Choragraph application has been optimally designed for visualising known and candidate proteins involved in membrane associations, membrane contact sites and dual-localised proteins. The paired colour settings and Organelle % filter can be particularly useful in searching for relevant proteins. Each protein has its own link to UniProt, so structures, extended annotation and further characteristics can be easily screened.

#### PM -ER

Setting the paired colour scheme to ER-PM efficiently highlights a string of proteins from the ER to the PM. Here we find well-known ER-PM contact proteins SYT1, 4 and 5, SYTF, PVA11 and 13 (Morello-López *et al*., 2026), calcium-dependent lipid binding protein CLB plus reticulon-like and HVA proteins involved in ER membrane curvature (Figure 7a). PDLP1 and a few members of the HIPP and MCTP families, known to be involved in plasmodesmata formation (Brault *et al*., 2019) and the ER quality control ERAD mechanism (Guo *et al*., 2021), are also present.

**Figure 7:**
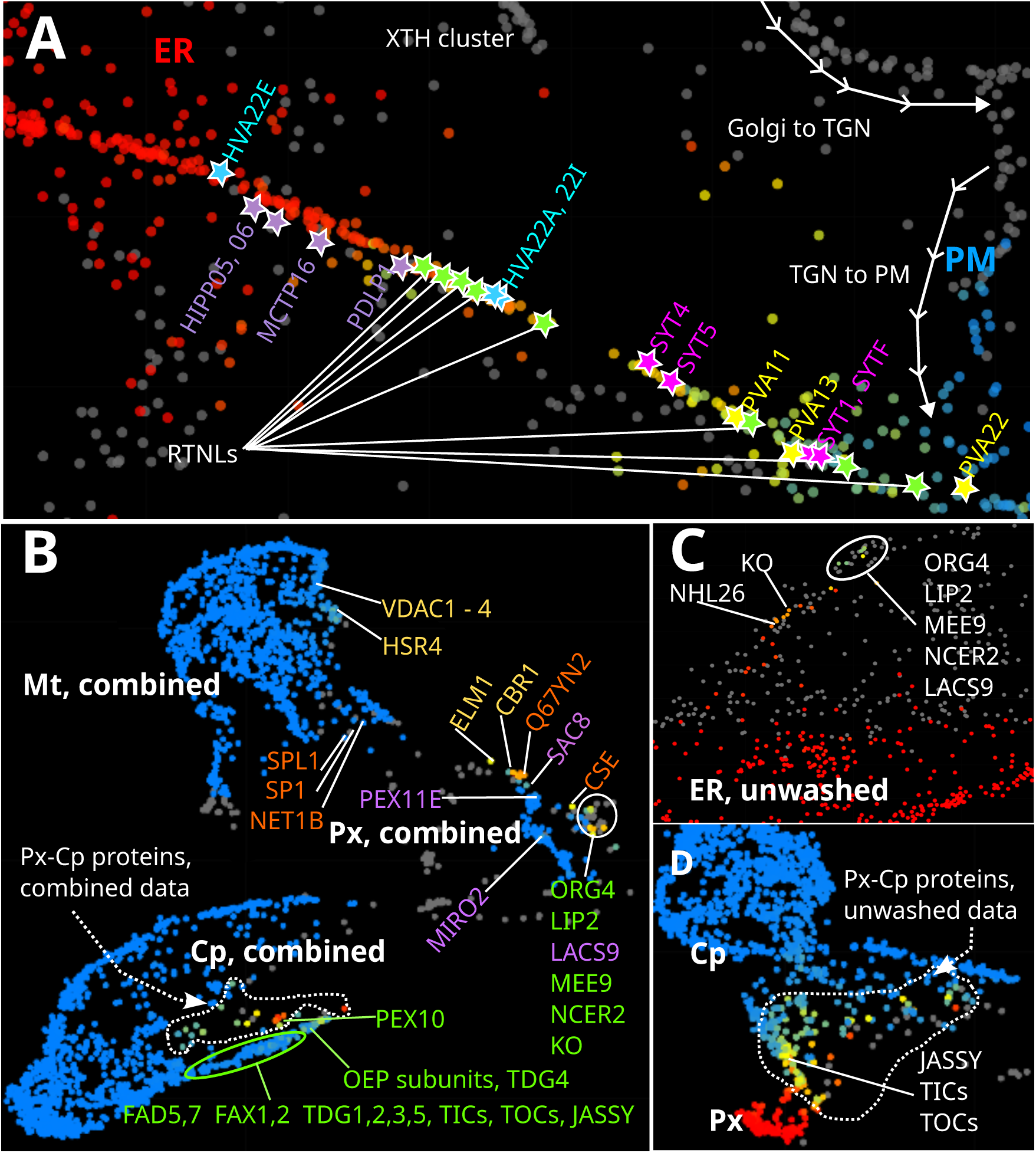
Membrane contact sites and metabolically linked proteins in the combined and unwashed datasets. a) Area of the combined dataset 2D plot showing the region of the ER-PM membrane contact site. RTNL proteins are marked as green stars and HVA22 proteins as blue. The established SYT and PVA protein markers for ER-PM MCSs are marked in magenta and yellow, respectively. Proteins more associated with the plasmodesmal ER-PM contact site are marked in purple. The background colour scheme is set to highlight the ‘ER-PM’ pair. Other features of this region are marked in white. b) Shared proteins between the ER and either the Mitochondria or Chloroplast or Peroxisome. Background colouring has been set so that intermediate-coloured proteins on the colour scale for all three organelle combinations show up together. Pair membership is shown by label colour: mainly ER-chloroplast proteins are labelled in green, mainly ER-mitochondrial in yellow, ER-peroxisomal in purple, peroxisomal-mitochondrial in orange. Proteins encircled by a solid white line are compared in Figure 7c. A group of chloroplast-peroxisomal proteins encircled by a dotted white are compared in Figure 7d. c) Locational comparison in the unwashed dataset with proteins encircled with a solid white line in the combined dataset (Figure 7b). d) Locational comparison in the unwashed dataset with proteins encircled with a dotted white line in the combined dataset (Figure 7b).

#### ER -Chloroplast/Peroxisome/Mitochondria

Glycerolipid biosynthesis occurs between the ER and Chloroplast and necessitates lipid transfer between the two organelles. The ER and chloroplastic locations of proteins involved in this shared pathway, such as trigalactosyltriacylglycerol desaturase (TGD), fatty acid desaturases (FADs), fatty acid exporter (FAX) etc. are reasonably well characterised (Hölzl and Dörmann, 2019). Lipid transfer proteins TGD4, and likely TGD5, have been previously identified at ER-Chloroplast MCSs (Xu *et al*., 2008), whilst LACS9 activates lipids for transfer (Huercano *et al*., 2025). If proteins are coloured according to sub-organelle markers (‘Other annotation’ in Choragraph), resolution of the outer envelope within the chloroplast cluster is evident. Using ER-Chloroplast organelle pair colouring highlights the end portion of this outer envelope extension. Along the extension we find several TGD proteins,TGD-like protein O80503, FAD6, FAX1 and 2. SQD2, which depends upon substrate delivery from the ER (Wang and Benning, 2012), is also present. TGD4 and a cluster of OEP complex subunits, which have distinct ER-Chloroplast profiles, define the very terminus of this extension (Figure 7b).

ABCD1 (PXA1), known to interact with ER tethering proteins (Esnay *et al*., 2020), localises to the ER facing edge of the peroxiome (Figure 7b), showing slight ER traits in its profile. Close by is SAC8, the orthologue of the yeast ER-PM membrane contact site protein sac1 (Despres *et al*., 2003), and which has a distinct ER-Peroxisome profile. The nearby ER-peroxisomal-mitochondrial GTPase activator protein Q67YN2 could plausibly be involved in MCS here, too. A subcluster to the right of the main peroxisome cluster, in the combined dataset, shows dual-localised proteins for chloroplast/peroxisome -ER/Golgi compartment combinations (solid white circle, Figure 7b). Caffeoyl shikimate esterase operates in the lignin biosynthesis pathway. This pathway depends upon precursors from the ER and Mitochondria and peroxisomal enzymes (Vanholme *et al*., 2010), so it is interesting to note the ER-peroxisomal-mitochondrial characteristics of this enzyme (Figure 7b and profile on webpage).

Using a sub-organellar colour scheme also shows a small cluster of mitochondrial outer envelope proteins at the ER-facing side of the mitochondrial cluster. Here we see HSR4 (Figure 7b), a protein that is functionally dependent on trafficking through ER-Mitochondrial MCSs. Close by we find mitochondrial outer membrane porins VDAC1, 2, 3 and 4 (Figure 7b). VDAC ion channels are known components of ER-Mitochondrial MCSs (Szabadkai *et al*., 2006). The mitochondrial fission protein ELM1 and the NADH-cytochrome b5 reductase CBR1, both involved in ER-mitochondrial MCSs, are found at the mitochondrial/peroxisome interface (Figure 7b) but have strong ER-mitochondrial profile traits.

Several proteins in this cluster, including LACS9 are, in the unwashed dataset, found in the ER-localised NHL26 cluster mentioned earlier, again suggesting this as a ER/Golgi to Chloroplast/Peroxisome/Mitochondria trafficking route (Figure 7c).

#### Chloroplast -Peroxisome - Mitochondria

The chloroplast, mitochondria and peroxisome are metabolically linked by pathways such as photorespiration. Highlighting the GO list of photorespiratory proteins (using Choragraph’s Protein ID box, then clicking “Select Filtered”) reveals that most of these are found between the peroxisome and mitochondria, except chloroplast contributors GLYK, PGLP1 and RUBISCO subunits.

Chloroplast-peroxisome connections seem to have been particularly sensitive to the addition of CO3-washed data to the combined dataset, as the TIC-TOC complex connection with the peroxisome is very clearly highlighted as dual-localised in the unwashed data (Figure 7d) but appears chloroplastic in the combined dataset (Figure 7c).The jasmonate biosynthesis protein JASSY has a clearly chloroplastic-peroxisomal profile in unwashed data, indicating how transient connections from compartment proximity can still be detected, dataset dependent. Likewise, photorespiratory proteins FtsHi5 and FtsHi2 (Wang *et al*., 2018) lose most of the peroxisomal component from their profiles with the CO3 wash, and dual-localised MIRO2 (Covill-Cooke *et al*., 2020) loses its mitochondrial profile traits, locating from the mitochondrial periphery to the centre of the peroxisome. PEX10 forms non-transient attachments at Peroxisome-Chloroplast contacts sites (Schumann *et al*., 2007). Here, in the combined dataset, it lies adjacent to the Outer Envelope chloroplastic protrusion that contains the Chloroplast-ER proteins (Figure 7c), with a clearly peroxisomal-chloroplastic profile.

SP1 and SPL1 are E3 ubiquitin ligases that function between all three compartments but localise preferentially to the peroxisome and mitochondria (Figure 7b), respectively, as has been demonstrated experimentally (Pan and Hu, 2018). Interestingly, the position (Figure 7b) and profile of actin-binding proteins NET1B suggests that it is involved in linking peroxisomes, mitochondria and possibly the ER, though it is generally associated with plasmodesmata (Deeks *et al*., 2012).

The 2D map projection does not differentiate between proteins that are dual-localised as a result of intermediate trafficking or those that end up at two locations. An example of the latter is a cluster that resembles a chloroplast-mitochondrial intermediate compartment that is composed mainly of aminoacyl tRNA synthetases, which are abundant in both organelles. Thus, mixing of organellar profiles from simultaneous residence creates this intermediate cluster effect. Strings of proteins extending out from the cytosol to the mitochondria and vice versa are, likewise, the product of dual targeted proteins. A prominent example here is DUT, where cytosolic, chloroplastic and mitochondrial spliceoforms seem to be present in our data, or SAL1 or NLP3, with chloroplastic and cytosolic spliceoforms.

### Using the Choragraph web application

The web application can be viewed at choragraph.org. The principal dataset from which most biological interpretations have been made is the combined dataset of 84 gradient samples and 9,438 proteins. Unwashed and CO3-washed datasets have also been included, however, as each contains unique proteins (9,850 and 9,970, respectively), that don’t meet the data sufficiency requirement for the combined set. Membrane structure is best preserved in the unwashed sample dataset but dual-localised protein profiles are often complicated by non-specific interactions. By combining datasets we try to balance non-specific protein interactions with preservation of membrane structures, as well as improving resolution with a larger dataset.

#### The 2D plot area

The webpage is divided into three main sections: the Protein Table, the 2D Data Plot and the Selected Protein area. The page initiates with the combined dataset. Hovering over data points shows Uniprot IDs, gene names and a brief description of each protein. Clicking on a datapoint shows the profile composition of this protein in the Selected Proteins area and selects the corresponding row in the Protein Table.

##### Datasets

Datasets are selected using the “Dataset” dropdown menu in the 2D plot area. The 2D Plot Area contains the 2-dimensional projection of multi-dimensional abundance profile data. The screen (x,y) coordinates of each point are derived from concatenated protein abundance profiles arising from density gradients: either 84 fraction samples for the combined profiles or 44 and 40 for the buffer- and CO3-washed datasets, respectively. When using the default UMAP projection method, proteins with similar overall profiles will end up with similar (x,y) coordinates, and so proteins residing in the same compartment will form clusters.

##### Projections

Each dataset can be viewed in its original, de-noised or latent or forms by selecting from the “Projection” dropdown menu. The original projection shows protein profiles where only the missing values have been filled by the neural network and existing values are unaffected. The de-noised projection shows protein profiles where the neural network has filled missing values and adjusted other values after learning from similar profile shapes. The latent projection shows proteins by their internal, latent representation within the DNN, which is then used for profile reconstruction/de-noising and compartmental classification. The relative position of clusters at a large scale in this projection is not related to the classification; this is merely the internal organisation of the DNN that has been optimised to support its predictive functions. In the original and reconstructed views a proximity between similarly classified profiles is preserved, so proteins that are related in biological data space are spatially close on the 2D projection. Switching between latent and de-noised UMAPs for the combined dataset, users can see how the Golgi is mostly isolated from the ER as a latent cluster; most of its profiles are distinct. However, in the denoised view the whole Golgi cluster is arranged close to the ER, and some Golgi profiles clearly are extending toward the ER.

As well as choosing between Original, De-noised and Latent data, users can choose between viewing the projection as a UMAP or PCA, and whether to view the entire subcellular dataset, or a map containing only proteins classified within the navigationally most complex parts of the cell. Unless otherwise stated, biological interpretations always refer to the combined UMAP of the entire subcellular proteome.

Finally, it should be generally noted that these maps are all low-dimensionality projections of higher-dimensionality data. As such, this involves compression and compromise in how points are arranged, and sometimes the less well connected groups are split in order to achieve the 2D representation. Also, some trails of protein spots may appear to pass through a cluster when, in a higher dimensional space, they would effectively be passing over the top.

##### Colour schemes

The “Colour by” dropdown menu presents options for colouring protein data points. Each colour scheme gives different insights into the data. These are summarised in the table below:

Selecting p1-value (1-class) as colours essentially illustrates the datapoints according to their single-ness. Consequently, highlighted areas reveal highways of dual-localised proteins. Regions highlighted by p2-value (2-class) colours show where protein profiles are not explained by either single- or dual-class assignments. This may reveal trafficking hubs, or intersections, represented by multi-localised proteins. Alternatively these may be proteins in a compartment not accounted for by the supervised training classes.

##### Other tools

The “Options” dropdown menu, next to the colour scheme, allows users to select data point sizes and opacity. Setting this to lower opacity and/or smaller spot sizes can be a useful means for assessing the density of clusters.

Hovering just below the “Dataset” and “Projection” dropdown menus highlights a row of tools for use in the plot area. Tool tip text appears over each one with a brief description, most of which are self-evident. The Box and Lasso selection tools allow users to subset regions of the plot, which can then be further filtered using the Protein Table. It is essential to click “Reset” after lassoing proteins, otherwise subsequent filters will be applied only to the subset.

#### The Protein Table area

For the selected dataset, the Protein Table lists proteins’ IDs, gene names, descriptions, locational predictions, annotated markers and profile completeness. By default the table and plot includes proteins with a completeness of 0.65 or better (i.e. up to 35% of the abundance profile was missing), which corresponds to the proteins used in the DNN training. The lower threshold can be adjusted down to 0.5, via the “Min, completeness” option, which will adjust the available proteins. A link icon at the left of each row takes users to the UniProt page for that protein, where full description, sequences and structures can be consulted.

##### Filtering the protein table

At the top left of the protein table are four dropdown menus. Clicking on “Protein ID” brings up a text entry box where users can paste in lists of candidate proteins either as uniport IDs, AGIs or gene names. Clicking on “Row Text” brings up three further menus. In these the users decides which column to filter, then selects the search parameters and finally inputs the search term. Note that the first option “IDs/description” searches the first four columns together and accepts input as IDs or keywords. The “Predicted Location” column uses text notation to show the probability with which classifications were made. Upper case text shows the most probable location(s) but will be prefixed with a question mark if this is not the majority location. Lower case text shows secondary locations and is suffixed with a question mark if there is no clear majority amongst secondary locations. It is essential to click “Reset” after the end of any filtering session, otherwise filters will be applied to previously selected proteins.

##### Downloading data from the table

At the top right are a series of Actions dropdown menus. “Select page” will highlight all proteins from that page on the 2D plot area. If proteins have been filtered already in the Search menus, “Select Page” will turn into “Select Filtered”. Note that this only happens if filter results have selected fewer than 200 proteins. Clicking “Download Data” brings up options for downloading the entire datasets, the current page or the filtered and selected proteins.

##### The Selected Protein area

The Selected Protein area contains all the information contained in the Protein Table for any one highlighted protein, with a pie chart showing all the compartment components identified with the protein’s profile. The same link icon as in the Protein Table takes users to the relevant Uniprot page.

## Summary and conclusions

Here we present a new subcellular spatial proteome of Arabidopsis, specifically targeted at investigating trafficking pathways and inter-organelle connections. By accurately reconstructing missing proteomic values, and classifying proteins against mixtures of known compartmental profiles, we have provided a high-resolution map of almost 10,000 proteins. We have made our results available on the Choragraph interactive web application, to provide easy access to our data and localisation predictions, so that membrane contact sites, trafficking pathways and dual-targeted proteins can be investigated.

By employing a context-aware deep learning approach with dual training objectives, we’re making extensive use of Arabidopsis hyperLOPIT data, which allows the number of proteins analysed to significantly exceed what has been achieved previously.

We’re making thousands of predictions of subcellular localisation in Arabidopsis, that extend well beyond what has been published to date, and in a manner that is coupled with an inherent measure of confidence. Also, many of the proteins that couldn’t be reliably assigned to a single compartment can now be explained as having varying degrees of dual-localisation.

The Choragraph application has enabled direct visualisation of trafficking pathways around the subcellular spatial proteome, with very high resolution of co-existing routes in high-traffic areas like the early secretory system. By analysing protein profile characteristics with regard to all possible compartment combinations, we can trace many of the physical associations made between a protein and different cellular membranes. We identify protein groups corresponding to current theories of trafficking and membrane contact, and demonstrate how Choragraph can be used to identify novel candidate proteins involved in membrane contact sites. We further demonstrate how our application can yield insights into lesser-understood routes such as Golgi-chloroplast trafficking, identify potential new compartments such as the XTH cluster, expansin/RALF cluster and even identify the trafficking machinery working on these clusters, such as the likely IRX12/laccase - Golgi interaction.

By presenting Choragraph as a web application, we aim to create a resource for hypothesis testing and validation that will benefit the wider plant cell biology research community.

## Data and code availability

Multi-localisation aware prediction results are available for our Arabidopsis datasets at: https://choragraph.org

This website allows for the display of classified proteomes as 2D UMAPs along with searchable tables of proteins and their classification scores etc.

All of the code to process the data, train the DNN models, make inference and generate SQLite3 output is available at: https://github.com/tjs23/choragraph

## Methods

### Preparation of biological material

Arabidopsis cell-suspension cultures were grown as described by (Parsons *et al*., 2012). Membranes were prepared as per the LOPIT density centrifugation protocol described in Parsons et al. 2019. After membrane pellets were harvested by centrifugation from gradient fractions, as described in Parsons et al. 2019, pellets were resuspended in either ice-cold standard 1X phosphate buffered saline, pH 7.4, or ice-cold 100mM NaCO3, pH11.0 for 30 min with occasional agitation. Pellets were recentrifuged at 100,000 x *g* for 30 min. Material was resuspended, digested, labelled and analysed according to Parsons et al. 2019.

### Training classifications

Lists of proteins with known subcellular localisation class, often referred to as “marker lists”, were largely derived from curated data that has been double-checked against literary references. For the Arabidopsis data presented here the markers come from Parsons et al. 2019. Ultimately these markers originate from a manual curation of Arabidopsis experimental protein localisation housed as SUBA (Hooper *et al*., 2017). This was the resource of choice, rather than UniProt, as it is easier to filter experimental data for potential dual- or multi-localised proteins at SUBA, which was of particular importance due to the nature of this study. The typical membrane-bound organelle classes were augmented with three extra classes; 1) for the ribosomes, 2) for 26S proteasomes and 3) for cellular cortex. Cellular cortex markers were retrospectively added after an initial iteration of Choragraph, in which it became apparent that coat protein complexes I and II were clustering together as a consequence of their attachment to the cytoskeleton. A UMAP of unprocessed input hyperLOPIT profiles shows that these extra classes account for clusters of proteomic profiles in regions that are otherwise not covered by organelle markers (Figure). This allows for more subtle discrimination of classes that might otherwise be largely interpreted as “cytosol” (to which they have some resemblance). Also, initial experiments where we did not include any markers in the regions resulted in other classes inappropriately bleeding into these clusters (by DNN, SVM and TAGM methods), when we are actually confident about what they represent.

Before the class lists were used in DNN training (as targets for a loss function) they were all subjected to a clean-up procedure to remove outlier examples that were inconsistent with the bulk of classification for their subcellular location (Supp Figure S2a). This was done in an automated manner by use of a simple k-nearest neighbours approach (Wilson, 1972). Accordingly, for a classified training protein, the 11 most highly correlated neighbour protein profiles that also carry a training classification were identified. Those target proteins where the majority of the neighbouring classifications did not match the target’s classification were then removed from the marker list (after all assessments were made). In large part, the marker pruning has the effect of inactivating markers that are most peripheral to their organelle. This means that when we train our DNN on class mixtures (see below) we are avoiding proteins that may be dual localised.

### Proteomic data preprocessing

Input data were normalised, on a per protein basis, by scaling each separate replicate LOPIT profile according to the sum of the protein’s values. Replicas were then concatenated into combined hyperLOPIT profiles. These were then scaled relative to the maximum of the whole, combined dataset, to fit all profiles within the range [0, 1], but without further per-protein, or per sample normalisation. Given that the per-protein normalisation of the replicate experiments represents a loss of absolute intensity information, we concatenated an extra value for each replicate experiment to the hyperLOPIT profiles (except in the causes where we were directly comparing to pre-imputed input). Here, each added value was the sum of the protein’s loge abundance values in the original replicate experiment, scaled per column to set the median value to 0.2.

The training profiles, and their corresponding target classifications, were then mixed to create the final input set. The test-train split of data, as described below, was done before profile mixing to keep the protein origins of the test set entirely separate. Profile mixing involved randomly selecting a fixed number for profiles (250 in this case) from each of two classes and combining these in random, but known, proportions to create mixed profiles, and corresponding target (class abundance) vectors. We excluded ribosomal and proteasomal classes from this mixing as they each represent one discrete, albeit large, multiprotein complex, rather than compartments containing many components. Overall, the mixing procedure acts to under- and over-sample the target training classes, dependent on size, and thus create a more balanced dataset.

More precisely, for a *M* × *N* sized data array *X* (one protein profile per row) and corresponding class identities *Y* (length M), a mixed profile 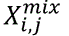 is created from randomly selected component profiles *X_i_* and *X_j_* with classifications *c* and *d*, according to:

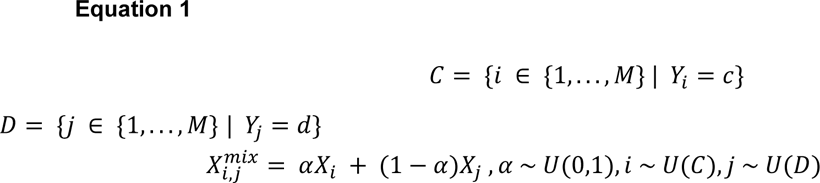

As *α* is chosen randomly within the range [0, 1] it thus provides a weighting for the addition of proportions of original profiles into a new, mixed profile. The corresponding target

classification vector for training, *Y^target^* with length K (one element for each class), was set as the vector of zeros for the unmixed classes and *α* or (1 − *α*) for the mixed classes.

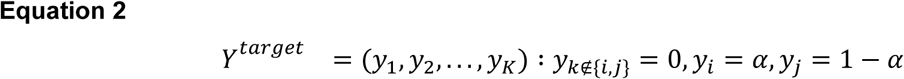

The paired profiles used in the mixing were set once, prior to training, for each model, but the *α* proportion was varied randomly for each pair each training cycle (epoch). To investigate predictive variance, caused by the test-train selection and profile pair choice, an ensemble of ten models, each with different mixed training pairs, was trained separately.

Each training run also included all the original, unmixed, single-origin proteomic profiles. Naturally these profiles corresponded to an organelle class weight of 1.0 on only one class.

The training data for each model was further augmented with an extra null class. This was created by randomly shuffling protein profile data within each column (i.e. columns never mix), with the condition that the random profiles do not have a Pearson correlation better than 0.5 to any real data point, whether its classification is known or otherwise. The size of the null set was set to be the same as the size of the original training dataset, i.e. not including mixed profiles. This null class acted as a decoy to define permissive boundaries of the known classes and to allow classifications to select “unknown” as a category where there is little evidence to choose any particular known subcellular localisation.

### Deep neural network architecture

The architecture of the deep neural network that we used to build the predictive models is illustrated in Figure 1b. Here input into the network was from both a minibatch of proteomic profiles and a reference consisting of all training proteomic profiles across the detected proteome. The reference Arabidopsis proteome input, where proteins had fewer than 35% missing values, contained 7,400 to 8,000 profiles, depending on the dataset.

Each input training batch is subject to random masking of 25% of values (set to zero) on-the-fly. The zeroed mask is also copied to the reference profiles corresponding to the same proteins as the input batch; to avoid direct copying of reference values.

Input profiles, containing the normalised proteomic data values (84, 44 or 40 here) are first compressed to a smaller vector with a single fully-connected layer operating on the last tensor dimension.

Normally distributed pseudorandom noise (*σ* = 0.05) and further weight dropout (10%) is then applied to the input batch alone, but not the proteome reference profiles. These provide regularisation and ensure that there are no exact matches between batch profiles and reference profiles.

Next there are several multi-headed cross-attention layers. Here the query is the input training batch and the keys are the reference profiles. Accordingly, information obtained from transforming and matching the batch queries to the reference keys allows the latent information in the query batch to be augmented in a context dependent manner. The number of attention layers, and the number of attention heads therein, was minimised to create a performant model that was as small as possible. This led to a final architecture with four cross attention layers, each with two attention heads. After the cross-attention layers, two separate outputs are generated. One of these is to predict subcellular localisation weights (see Equation 2) and the other to reconstruct the masked input values.

The reconstruction of the proteomic profiles from the cross-attention vector output is done with two layers of fully-connected Bayesian variational inference (BVI) layers, as made available in Tensorflow as DenseReparameterization (Kingma, Salimans and Welling, 2015; Kingma and Welling, 2022). The first of these uses a GELU activation function for non-linearity and the second expands the latent vector to the original hyperLOPIT profile size and attenuates with sigmoidal activation.

During training, the optimised network parameters of the BVI layers correspond to both the mean of the network weights and their normally distributed variance. The purpose of the BVI layers is to create a model with stochastic output during inference. Network weights may be sampled according to their fitted, normal distributions, and thus the model output is sampled to generate distributions of predictions. In turn these may then be used to estimate output dispersion, and p-values etc.

The prediction of subcellular class, or proportions thereof, also uses BVI layers. Here three fully-connected layers, with intervening GELU non-linear activations, are used to shrink the latent vector size from the cross-attention representation to per-class subcellular location scores (effectively proportions).

### Deep neural network training

The test-train input partitioning was done by shuffling proteins randomly into 10 sets and then selecting each of those sets in turn to act as test data and combining the other nine sets to be training data. Training data order was randomised each training epoch, but (as mentioned above) data null-class augmentation and profile mixing was only done once at the start.

An ensemble of ten independently trained models were then trained, one for each partition, and used for future inference. In all cases, optimization of DNN weights was done using the ADAM optimizer (Kingma and Ba, 2017) with a cosine annealing schedule (Loshchilov and Hutter, 2017) to set a diminishing learning rate that range from 10^−3^ to 10^−5^ over 75 training epochs. Overall, tracking of the test and training loss and accuracy showed that convergence was good with no evidence of overtraining.

The loss function for training was the summation of a categorical output loss and a masked-input profile reconstruction loss. The reconstruction loss was mean absolute error; comparing unmasked original profiles to reconstructed output. The categorisation loss was the negative log-likelihood of the observed localisation proportions given a 2-trial multinomial distribution, based upon the 2-class proportions of the training subcellular localisation mixtures. Details of the loss functions are described below.

### Deep neural network inference

Inference was done for the whole proteome dataset, where sufficient data exists. For the analyses we present here this was judged to be <35% missing points. However for the Choragraph web application we relax this to <50% missing points, so that users may investigate borderline situations. Inference consisted of sampling 100 paired outputs from each of the ensemble of ten separately trained models. Thus, 1000 class vectors (containing proportions/scores) and 1000 reconstructed profiles were sampled as output for each protein.

Given these outputs, a heuristic was used for delineating likely single- and dual-localised proteins. In reality there is a continuum of predictions with associated errors. Hence, as described below, we also sought to assign confidence by estimating the single-class prediction p1-value, and the dual-class prediction p2-value.

A protein’s classification was deemed to be singular, i.e. corresponding to only one subcellular localisation class, if the p1-value is less than 0.001 and the second highest scoring class has a mean class score (proportion) less than 0.2. Otherwise, a protein was deemed to be likely dual localised if the top two classes (with the two greatest average scores) have a combined p2-value of less than 0.001.

Proteins which do not fit either of these conditions were left as uncertain in classification. In a few cases these may be discernable as having more than two sensible subcellular localisations. However, these are generally interpreted as undetermined as they cannot be easily explained by the two-class models, especially where there is notable weight contribution from the null class. Overall, classes which contribute to less than 0.05 average proportion were not noted, though these values are available in the full inference data.

Where standardized measures of singleness and dualness were required these were calculated by considering the proportion of probability density (after fitting to the score distributions) at or above a score threshold of 0.8. The “singleness” measure considered the proportion of probability density for the highest scoring classes (by ensemble mean). The “dualness” measure considered the combined proportions for the top-two scoring classes, minus the singleness measure; i.e. the dualness is the gain above singleness.

### P-value estimation

The 100 class proportions (predicted alpha values) from each model were fit to a beta distribution using the .fit() method of scipy.stats.beta (Virtanen *et al*., 2020). Accordingly, sampled values from each model were fit to a smooth distribution bounded in region [0,1]. The ten fitted beta probability density functions, one for each model, were then averaged to generate a smooth distribution representing the combined ensemble’s 1000 points. This mean-of-betas representation was then used in the estimation of p-values for a given single-or dual-location classification.

To estimate classification p-values for each subcellular localisation we considered the null hypothesis that the sample is actually output from any class other than the one(s) being evaluated. The p-value for a protein having a given class is estimated as the probability of obtaining an output value at or above the class’s 10th percentile (critical) value, under the assumption that the null hypothesis is true. This critical value was set heuristically to indicate when compartmental mixing becomes biologically interesting. Thus, the p-value is calculated as the sum of tail integrals from the other classes, and these tail integrals are estimated from the fitted mean-of-betas.

The critical value *ck* used for class *k* is the 10% percentile (*P*0.1) output proportion for that class *ŷ_k_* over all *S* inference samples :

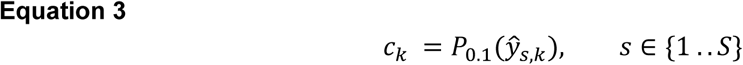

For a total of *M* models and *K* classes, with beta distribution parameters *αi*,*m* and *βi*,*m* fitted to the outputs of model *m* at class indices *i*, the p-value for class *k* is:

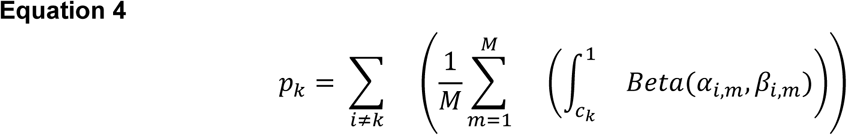

To extend the p-value calculation to proteins that are dual localised the two most likely classes are combined by summing their output scores. The calculation then proceeds as above with the exception that the null hypothesis now relates to all but the chosen two classes. Accordingly, a roughly 50%:50% dual localised protein can have a single-class p-value near 0.5 but a dual-class p-value near 0.0. We refer to the single-class and dual-class hypothesis p-values as the p1-value and p2-value respectively.

### Missing value reconstruction

The reconstructed proteomic profiles that were output from the model inference (along with class proportions) were used to create complete without missing values datasets. These full representations may then be analysed further or visualised in 2D by PCA and UMAP etc. The ensemble of DNN models yielded 1,000 predictions for each datapoint in each protein profile. We took the median of the points at each profile position to estimate the best, denoised profile value.

We make available two versions of a reconstructed dataset, one is a fully synthetic dataset that was output from the DNN for each protein. The second is a hybrid of the original input data and reconstructed DNN output, where the DNN output is only used to replace the original missing/zero values. While both datasets are similar the latter naturally is a closer representation of the recorded proteomic data.

### Loss functions

The loss for reconstruction of masked proteomics profiles, of length *N*, was calculated as the mean absolute error between output prediction *ŷ* and target *y* where the target elements are non-zero:

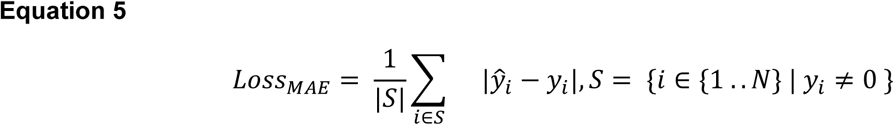

The loss for the prediction of the categorical proportions (an array of length *K*) was calculated as the negative log-likelihood from a 2-trial multinomial distribution, using the TensorFlow probability (Dillon *et al*., 2017) class tfp.distributions.Multinomial. With multinomial coefficient *C*, the loss may be expressed as:

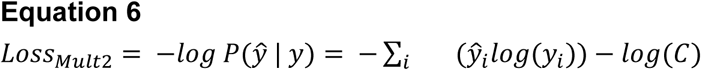

### Prediction metrics

Predictive accuracy was calculated as the proportion of all predictions that were correct;

*Accuracy* = *n*(*correct*) / *n*(*predictions*). For proteomic profile reconstruction, accuracy was calculated with a threshold of 0.1 on the normalised scale; a point was deemed to be sufficiently close to be correct if the absolute difference between predicted and target value was at or within this threshold. Also, reconstruction accuracy was only evaluated for the originally non-zero profile points, whether masked or unmasked.

Class precision and recall were calculated from the true positive (TP) false positive (FP) and false negative (FN) class predictions within a partitioned test data set as *Precision* = *TP*/ (*TP* + *FP*) and *Recall* = *TP*/(*TP* + *FN*). Here “positive” corresponds to predictions of the specified class and “negative” corresponds to predictions for any other classes.

The class-wise F1 score is then readily calculated from the class precision and recall as *F*1*class* = 2 ⋅ *Precision* ⋅ *Recall*/(*Precision* + *Recall*). The macro F1 was then taken as the arithmetic mean of all of the class-wise F1 scores.

### UMAPs

Two-dimensional UMAP projections were calculated using umap-learn (https://umap-learn.readthedocs.io) (McInnes, Healy and Melville, 2020). Throughout, the UMAP parameters used were: *num. neighbours; **30**, minimum distance; **0.1**, distance metric; **‘correlation’***.

### SVM training

The SVM classification models were fitted to exactly the same test-train data splits and class markers as the DNN models. Accordingly, an ensemble of 10 SVM models was trained, with the sklearn.svm.SVC classifier (Chang and Lin, 2011; Pedregosa *et al*., 2011), available within the scikit-learn Python module (https://scikit-learn.org) and probability scores were generated via sklearn.svm.SVC.predict_proba(). A mean score, over all models, corresponding to 0.8 or more was used to select the confident SVM predictions, and this threshold closely matched the confident predictions from the equivalent DNN-based models.

### TAGM

We ran all data-sufficient abundance profiles from the unwashed dataset through the TAGM MCMC prediction pipeline (Crook *et al*., 2018), with the same culled training markers used by the DNN. Training was run with the following parameters: *numIter;**16000**, burnin; **4000**, thin; **20**, numChains; **8***.

