## Supplemental Figures S1 - S5 for "Choragraph: A deep-learning approach for the analysis of spatial proteomics reveals subcellular Arabidopsis protein trafficking routes and multi-residency"

### Supplemental Figure 1

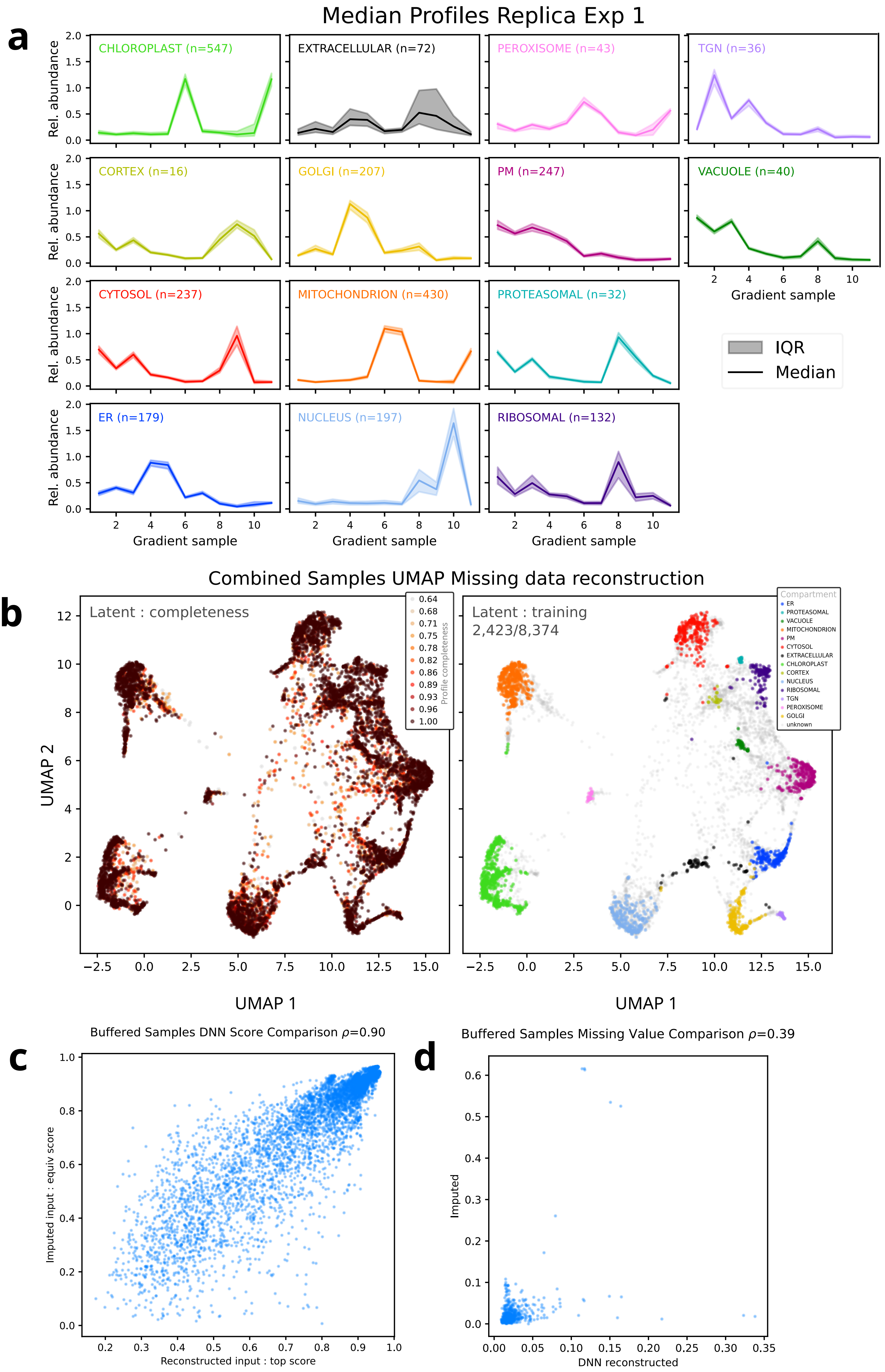

### Supplemental Figure 2

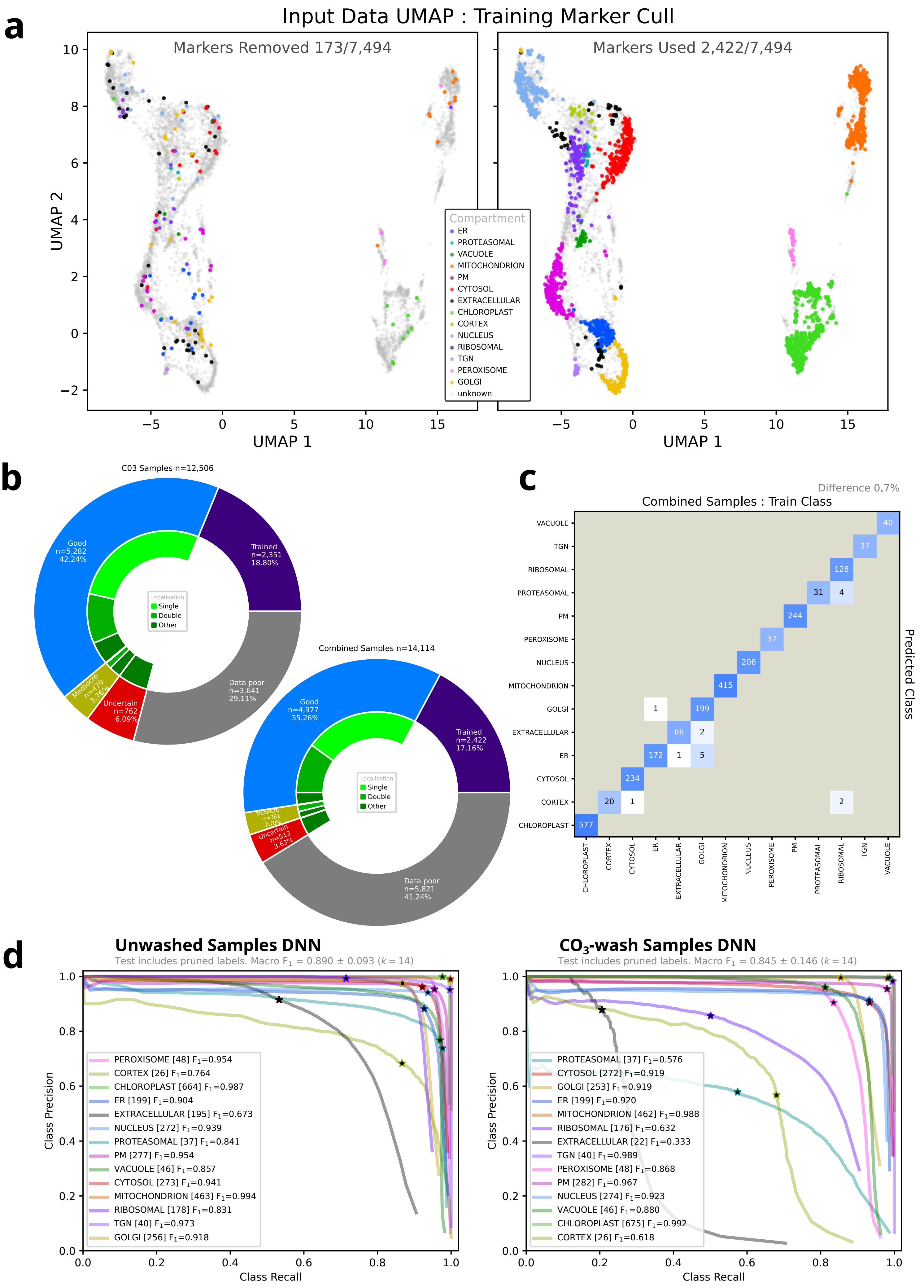

### Supplemental Figure 3

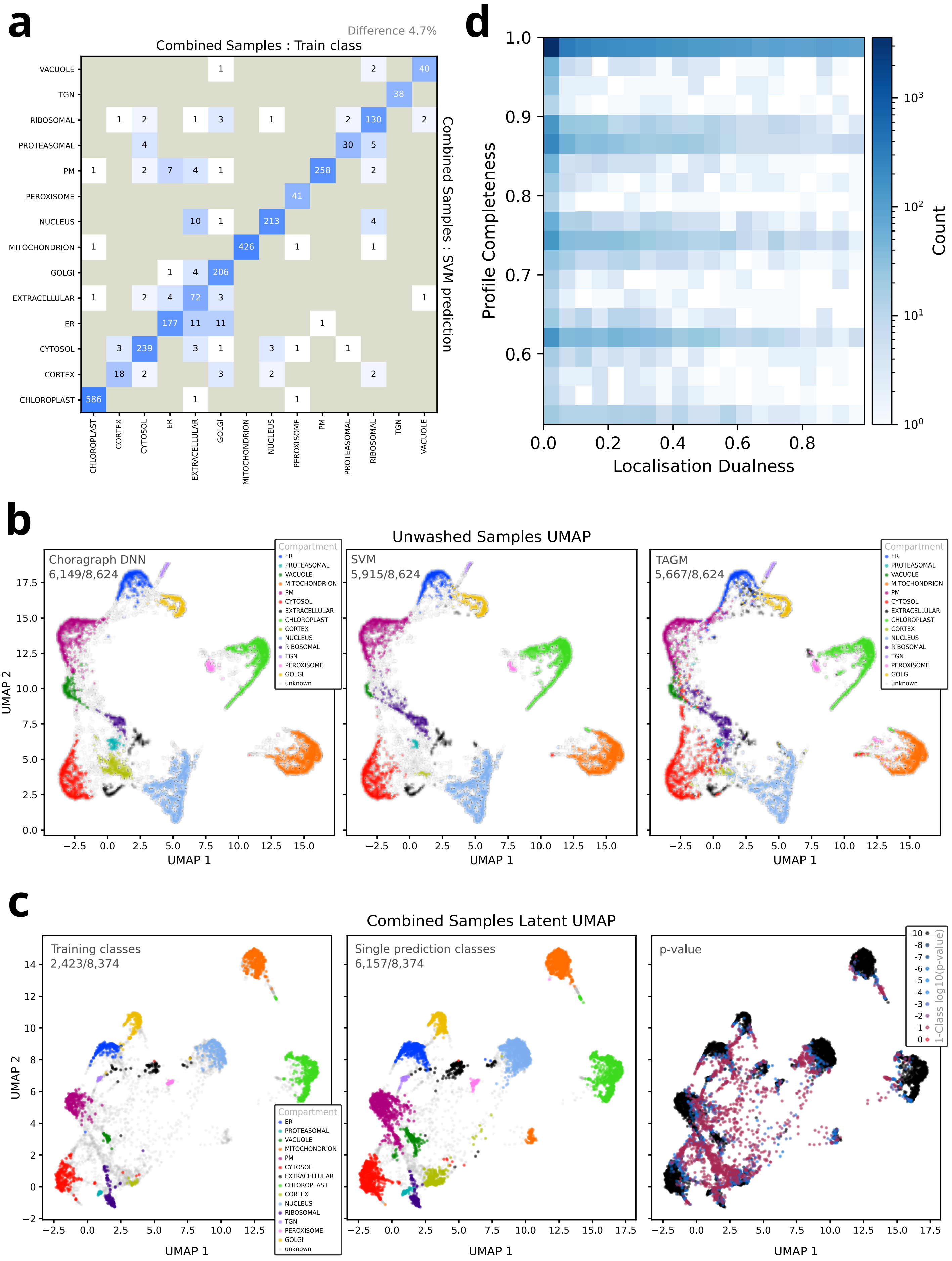

### Supplemental Figure 4

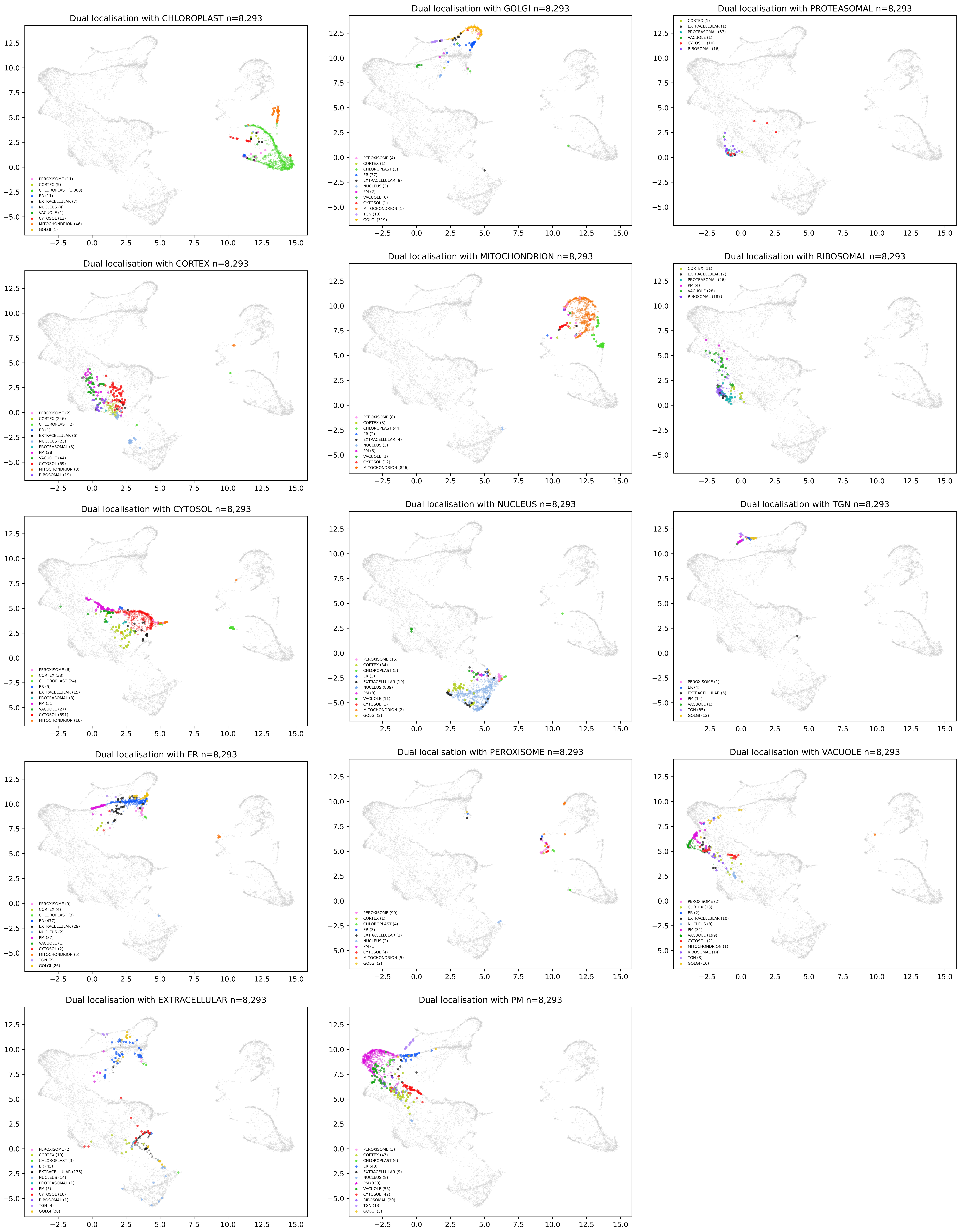

### Supplemental Figure 5

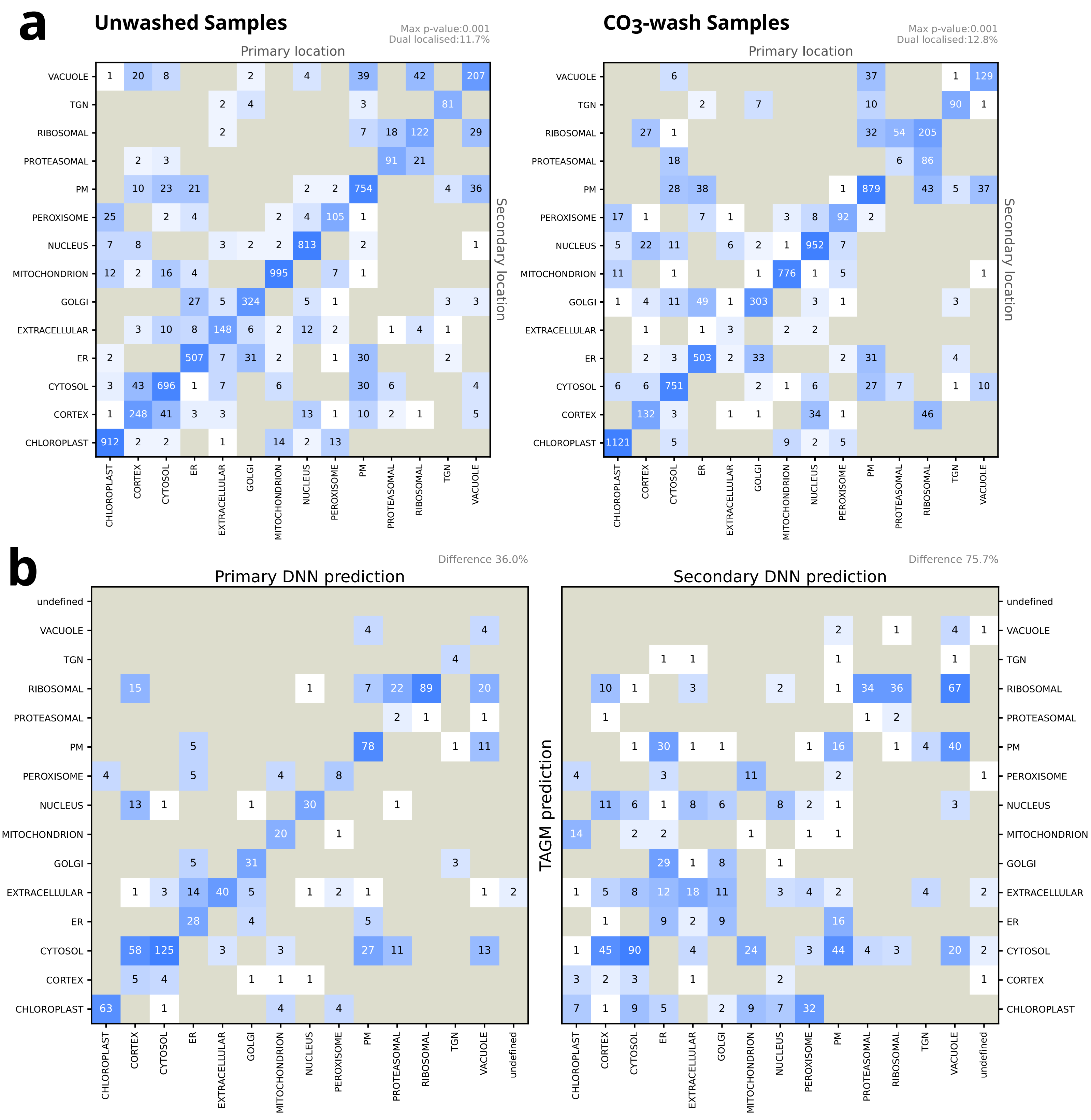

Supp. Figure S1:

a) Average proteomic profiles for the various subcellular localisation classes used in DNN training. Dark lines indicate the median marker profile for that class, and the shaded areas indicate the interquartile range. Data are shown for the first replica buffer wash samples.
b) 2D UMAP projection of the DNN latent vectors. Each point represents a different protein and is coloured according to the fraction of values missing in the original profile (left) or according to the subcellular compartment classification used in training (right). The data shown is for the combined dataset.
c) The correspondence between DNNs top-class classification scores (horizontal axis) and the equivalent scores (for the same classes) when the input is pre-filled using KNN imputation (vertical axis).
d) Comparison between pre-filling missing values using KNN imputation and DNN based on-the-fly reconstruction of values.

Supp. Figure S2:

a) 2D UMAP projections of input proteomic profiles coloured according to known, marker classification, split according to whether the marker was considered a single-class outlier and disregarded (left) or kept for training with the DNN classifier (right).
b) Overview of how protein abundance profiles were used with the DNN classification, for the combined LOPIT data; 8 replicas over two conditions (left) and the carbonate wash data; 4 replicas (right). The outer ring illustrates the proportions of profiles used as training markers, excluded from classification due to missing values (>35%) or classified with various degrees of confidence, as given by p-value. The inner green ring shows the proportions of the classified profiles that were considered, single-, dual- or multi-localised.
c) Confusion matrix to show the tally of differences between the known marker classification for test profiles, which were excluded from model training (columns) and their predicted DNN class (rows), with p-value < 0.001. Results are aggregate counts over 10 separately trained DNN models, where each marker protein was used in a test set once. Diagonal elements indicate identical classification and off-diagonal elements indicate differences.
d) Precision-recall curves and maximum F1 scores for each of the singly classified subcellular location classes using DNN models trained on the unwashed dataset (left) or carbonate-wash dataset (right). Data shown are averages of ten independent test-train data splits, and the same data splits were used for both DNN and SVM. In all cases the test proteins included markers that were pruned, and never used in training.

Supp. Figure S3:

a) Confusion matrix for the training of the SVM models. The tally of differences between the known marker classification used in training (columns) and the SWM predicted class using probability threshold >= 0.8 (rows). Results are aggregate counts over 10 separately trained SVM models, where each marker protein was used in a test set once.

b) 2D UMAP projections of DNN latent vectors, for each protein, coloured according to training marker classification (left), predicted single-class classification, with $p_1$-value < 0.001 (middle) and single-class $p_1$-value (right). The DNN latent vector is the internal representation of the model before the split between classification and profile reconstruction layers.

c) 2D UMAP projections of denoised proteomic profiles coloured according to predicted, singular subcellular class for different computational methods. Classification is shown for the DNN with $p_1 < 0.001$ (left) SVN with probability >= 0.8 (middle) and TAGM with MCMC probability >= 0.9999 (right).

d) A 2D density histogram to show any relationship between the degree missing/present profile and the dual-localisation of a protein; the gain in the fraction of DNN scores > 0.8 when combining primary and secondary classes. Here the notable horizontal pattern comes from when a protein is missing from one of the replicate experiments.

Supp. Figure S4:

An array of panels with the same 2D UMAP projection, but highlighted differently to show the single- and dual-localised proteins relevant for each subcellular location considered. In each panel, the single-compartment proteins for the reference class are indicated with a star. Dual-localised proteins are indicated with circles and coloured according to the other, non-reference component of the classification. Data is shown for the denoised proteomic profiles from the complete dataset.

Supp. Figure S5:

a) Comparison matrices to illustrate the DNN assignment of proteins to one or two subcellular classes using the data for the unwashed samples (left) and carbonate-wash samples (right). Diagonal elements indicate assignment to a single-class ($p_1 < 0.001$). Off-diagonal elements indicate assignment to dual classes ($p_1 > 0.001$, $p_2 < 0.001$), as specified by column (primary class) and row (secondary class).

b) Comparison matrices to tally how dual-localised proteins identified by the DNN are classified by the TAGM MCMC method. Diagonal elements indicate identical classification. The matrices on the left refer to the primary DNN classification (with the largest proportion) and the matrices on the right refer to the secondary DNN classification. Classification thresholds for the different methods were: DNN $p_1 > 0.001$, $p_2 < 0.0001$; TAGM MCMC probability >= 0.9999.
